# Cristae undergo continuous cycles of fusion and fission in a MICOS-dependent manner

**DOI:** 10.1101/654541

**Authors:** Arun Kumar Kondadi, Ruchika Anand, Sebastian Hänsch, Jennifer Urbach, Thomas Zobel, Dane M. Wolf, Mayuko Segawa, Marc Liesa, Orian S. Shirihai, Stefanie Weidtkamp-Peters, Andreas S. Reichert

**Affiliations:** Institute of Biochemistry and Molecular Biology I, Medical Faculty, Heinrich Heine University Düsseldorf, 40225 Düsseldorf, Germany; Center for Advanced Imaging, Faculty of Mathematics and Natural Sciences, Heinrich Heine University Düsseldorf, 40225 Düsseldorf, Germany; Department of Medicine, Nutrition and Metabolism Section, Evans Biomedical Research Center, Boston University School of Medicine, 650 Albany Street, Boston, MA 02118, USA; Department of Medicine, Division of Endocrinology, and Department of Molecular and Medical Pharmacology, David Geffen School of Medicine, University of California Los Angeles, CA 90095, USA

## Abstract

The mitochondrial inner membrane can reshape under different physiological conditions. How and at which frequency this occurs *in vivo* and what are the molecular players involved is unknown. Here we show using state-of-the-art live-cell stimulated emission depletion (STED) super-resolution nanoscopy that crista junctions (CJs) are dynamically fusing and dividing in a reversible and balanced manner at a timescale of seconds. CJ dynamics is strongly reduced in the absence of the MICOS subunit MIC13. Staining of the cristae membrane using different protein markers or two inner mitochondrial membrane-specific dyes revealed that cristae also undergo continuous cycles of fusion and fission. These processes are dependent on MIC13 and occur at a timescale of seconds, resembling CJ dynamics. Our data further suggest that MIC60 acts as a docking platform pioneering CJ formation. Overall, by employing a variety of advanced imaging techniques including FRAP (Fluorescence-Recovery-After Photobleaching), SPT (Single-Particle-Tracking), live-cell STED and confocal Airyscan microscopy we demonstrate that cristae undergo continuous cycles of fusion and fission in a manner that is mechanistically linked to CJ formation and dynamics.

## Introduction

Mitochondria are vital organelles with key roles in energetics and metabolism of the cell. The ultrastructural morphology of this double-membrane enclosed organelle is highly variable and altered in numerous human disorders (Zick et al, 2009). The internal mitochondrial structure is characterized by invagination of the inner membrane (IM) called cristae. The IM that closely remains apposed to the outer membrane (OM) is called the inner boundary membrane (IBM). The cristae membrane (CM) connects the IBM via a highly curved, circular or slit- or pore-like structures called crista junctions (CJs) (Acehan et al, 2007; Perkins et al, 1997; Perkins et al, 2003; Perkins et al, 2010; Rabl et al, 2009; Zick et al, 2009). CJs are highly conserved with a diameter of 12-40 nm and were proposed to act as diffusion barriers for proteins or metabolites (Frey et al, 2002; Mannella et al, 2013; Mannella et al, 1994; Mannella et al, 2001). Thus, the presence of CJs could create several mitochondrial subcompartments by separating IBM from CM and intermembrane space (IMS) from intracristal space (ICS). Indeed, the CM is enriched in proteins involved in oxidative phosphorylation (OXPHOS), mitochondrial protein synthesis or iron-sulfur cluster biogenesis, whereas the IBM mainly contains proteins involved in mitochondrial fusion and protein import (Vogel et al, 2006). CJs could regulate bioenergetics by limiting the diffusion of ADP/ATP and affect the pH gradient across the IM (Frey et al, 2002; Mannella et al, 2013; Mannella et al, 1994; Mannella et al, 2001). As early as 1966, isolated mitochondria are known to occur in different morphological states, condensed (matrix condensed, high ADP, state III) or an orthodox (matrix expanded, low ADP, state IV) state, depending on the bioenergetic status (Hackenbrock, 1966; Hackenbrock, 1968; Hackenbrock, 1972). Later, tomographic images of mitochondria undergoing this transition indicated that remodelling of the IM occurs in isolated mitochondria (Mannella et al, 1994). Cristae exist in different shapes and sizes depending on the physiological, bioenergetic or developmental cues (Mannella et al, 1994; Nicastro et al, 2000; Zick et al, 2009). Moreover, the general ability of cristae or CJs to dynamically remodel is well exemplified during apoptosis, where widening of CJs is observed, promoting cytochrome *c* release from the intra-cristal space into the cytosol (Frezza et al, 2006; Scorrano et al, 2002). However, molecular mechanisms for cristae and CJs remodelling in response to metabolic and physiological adaptations are not known. Aberrant and altered cristae are associated with several human diseases including neurodegeneration, cancer, diabetes and cardiomyopathies (Vincent et al, 2016; Zick et al, 2009) but their relevance to disease progression is unclear.

The formation of a highly curved, membrane-embedded CJ structure is likely to require an intricate partnership between phospholipids and scaffolding proteins (Koob & Reichert, 2014; Quintana-Cabrera et al, 2017; Rampelt et al, 2016). We identified that Fcj1(*formation of crista junction protein 1*)/Mic60 resides preferentially at CJs in yeast and its deletion leads to complete loss of CJs with cristae arranged as concentric stacks separate from the IBM. In addition, Fcj1/Mic60 and Su e/g (subunits of F_1_F_o_-ATP synthase) act antagonistically to control F_1_F_o_-ATP synthase oligomerization and thereby modulate formation of CJs and cristae tips (Rabl et al, 2009). Several groups have identified a large oligomeric complex termed MICOS (Mitochondrial contact site and cristae organizing system) which is required for the formation and maintenance of CJs and contact sites between IM and OM (Harner et al, 2011; Hoppins et al, 2011; von der Malsburg et al, 2011). The MICOS complex contains at least seven subunits in mammals: MIC10, MIC13, MIC19, MIC25, MIC26, MIC27 and MIC60 named after a uniform nomenclature (Pfanner et al, 2014). MIC60 and MIC10 are considered to be the core components of the MICOS complex in baker’s yeast as their deletion causes complete loss of CJs. MIC60 has a binding interface for a variety of proteins including TOM complex, OPA1, SAM/TOB, Ugo1 (mammalian homolog SLC25A46), DnaJC11, CHCHD10, and DISC1 (Disrupted-in-Schizophrenia 1) and is proposed to provide the scaffold for MICOS as well as contact between IM and OM (Abrams et al, 2015; Barrera et al, 2016; Genin et al, 2015; Glytsou et al, 2016; Körner et al, 2012; Ott et al, 2012; Pinero-Martos et al, 2016; Xie et al, 2007). MIC10 contains conserved glycine motifs in its transmembrane domain that are crucial for MIC10 self-oligomerization, and are required for the stability of CJs (Barbot et al, 2015; Bohnert et al, 2015; Milenkovic & Larsson, 2015). MIC10 additionally interacts with the dimeric F_1_F_o_-ATP synthase and promotes its oligomerization (Eydt et al, 2017; Rampelt et al, 2017). Both MIC60 and MIC10 have the capability to bend membranes (Barbot et al, 2015; Hessenberger et al, 2017; Tarasenko et al, 2017). Using complexome profiling, we identified MIC26/APOO and MIC27/APOOL as *bona fide* subunits of the MICOS complex (Koob et al, 2015; Weber et al, 2013). Depletion or overexpression of MIC26 or MIC27 lead to altered cristae morphology and reduced respiration. MIC27 binds to cardiolipin, the signature lipid in mitochondria (Weber et al, 2013). The non-glycosylated form of MIC26 is a subunit of the MICOS complex, but not the glycosylated form (Koob et al, 2015). Recently, another group and we have discovered that MIC13/QIL1 is an essential component of the MICOS complex and is responsible for the formation of CJs (Anand et al, 2016; Guarani et al, 2015a). Loss of MIC13 resulted in reduced levels of MIC10, MIC26 and MIC27, accompanied by impaired OXPHOS. The protein levels of MIC60, MIC19 and MIC25 remain unaltered, suggesting that MICOS comprises two subcomplexes, MIC60/25/19 and MIC10/13/26/27 with MIC13 acting as a bridge between both subcomplexes (Anand et al, 2016; Guarani et al, 2015a). Altered levels of MICOS components and their interactors are associated with many human diseases such as epilepsy, Down syndrome, frontotemporal dementia-amyotrophic lateral sclerosis, optic atrophy, Parkinson’s disease, diabetes, and cardiomyopathy (Abrams et al, 2015; Penttila et al, 2015; Zerbes et al, 2012). Mutations in *MIC60* have been found in Parkinson’s disease (Tsai et al, 2018). Mutations in *MIC13/QIL1* lead to mitochondrial encephalopathy and hepatic dysfunction (Godiker et al, 2018; Guarani et al, 2016; Russell et al, 2019; Zeharia et al, 2016).

Here we studied cristae membrane remodelling in living cells and what role the MICOS complex plays in this context. To study systematically intra-mitochondrial dynamics of CJs and cristae, we devised a novel state-of-the-art method of live-cell STED super-resolution nanoscopy using the C-terminally SNAP-tagged versions of distinct mitochondrial proteins marking CJs and cristae. MIC10- and MIC60-SNAP marking CJs move dynamically and join and split continuously at a time-scale of seconds in a MIC13-dependent manner. In conjunction, we also observed that cristae marked by ATP5I-SNAP and COX8A-SNAP or by IM-specific dyes undergo continuous fusion and fission cycles in a balanced manner in a similar timescale of seconds. Overall, by improving temporal and spatial resolution using live-cell STED super-resolution nanoscopy in combination with the SNAP-tag technology and use of newly generated genetic models lacking MICOS subunits we achieved to resolve cristae membrane dynamics. Based on these findings we propose a model rationalizing the physiological importance of the MICOS complex and of cristae remodelling.

## Results

### Mammalian MIC10 and MIC60 are required for cristae morphogenesis and cellular respiration

MIC60 and MIC10 are the core subunits of the MICOS complex that are also evolutionary well conserved (Huynen et al, 2016; Munoz-Gomez et al, 2015). As the mammalian MICOS complex is not well understood, we obtained human *MIC10* and *MIC60* knockout (KO) HAP1 cells. *MIC10* KO and *MIC60* KO have 29 bp deletion in exon 1 and 10 bp deletion in exon 8, respectively, leading to frameshift and subsequent loss of the respective proteins (Fig 1B). Analysis of electron micrographs from these cells revealed that CJs are virtually absent in *MIC10* and *MIC60* KO cells (Fig 1A, 1C). The cristae membrane (CM) appears as concentric rings detached from the IBM (Fig 1A), consistent with earlier observations in baker’s yeast and mammalian cells (Callegari et al, 2019; Harner et al, 2011; Hoppins et al, 2011; John et al, 2005; Rabl et al, 2009; von der Malsburg et al, 2011). In addition, the abundance of cristae per mitochondrion is reduced (Fig 1D). While the knockout of *MIC10* primarily causes a selective destabilization of the MICOS subcomplex comprising MIC13, MIC26 and MIC27, loss of MIC60 results in a clear destabilization of all subunits of the MICOS complex (Fig 1B), confirming its role as a main scaffolding subunit of MICOS. The basal and the maximal oxygen consumption rates of *MIC10* and *MIC60* KOs are significantly decreased compared to controls (Fig 1E, IF), confirming the role of CJs in ensuring full bioenergetic capacity.

**Figure 1.**
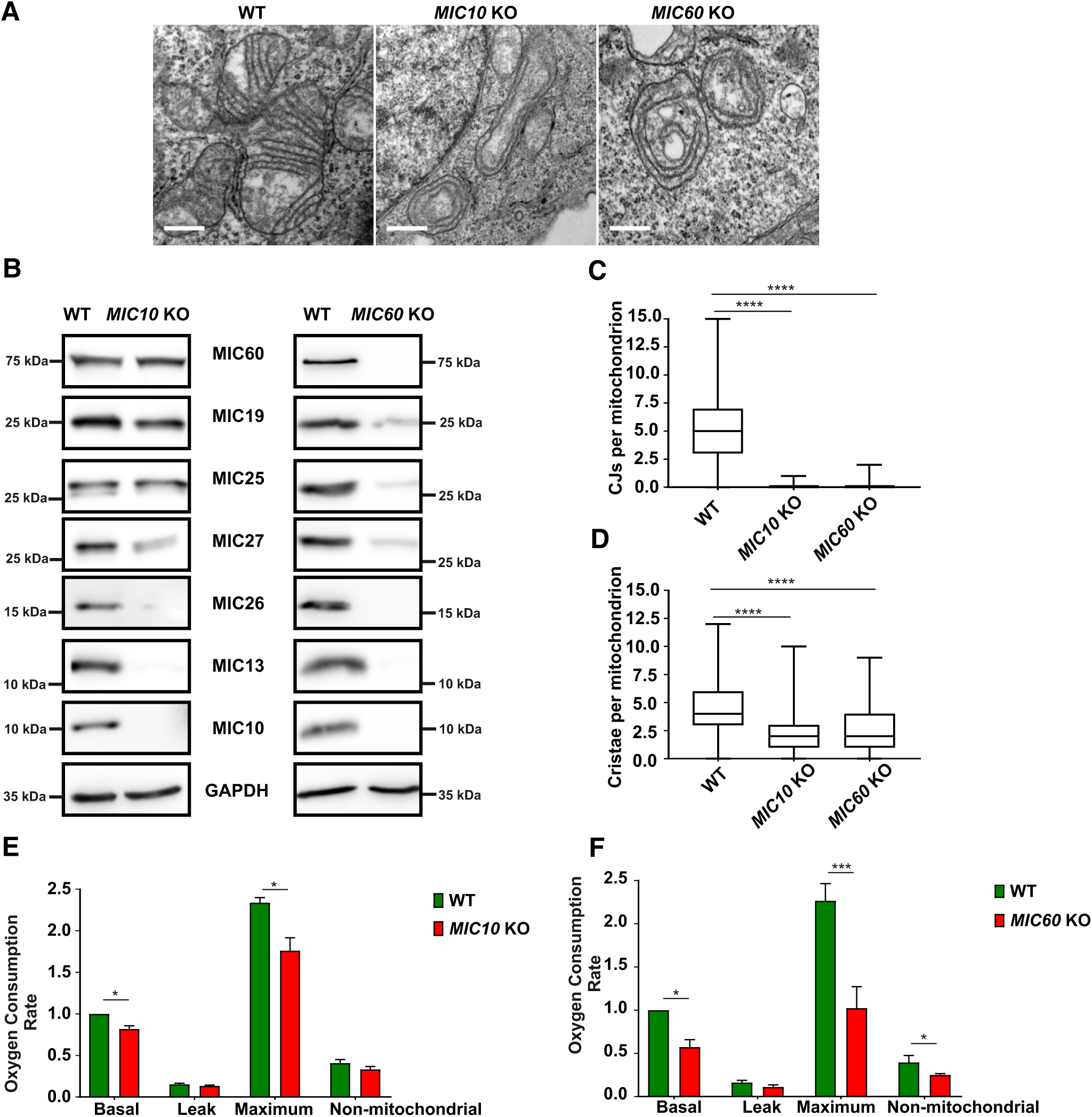
*MIC10* and *MIC60* KO HAP1 cells show loss of crista junctions and impaired cellular respiration. **A** Representative electron micrographs of WT, *MIC10* KO and *MIC60* KO HAP1 cells. Scale bar 0.5 µm. **B** Western blot analysis of lysates from WT and *MIC10* KO or *MIC60* KO HAP1 cells. *MIC10* KO show a drastic reduction of MIC13, MIC26 and MIC27 protein levels while protein levels of other MICOS components remain unchanged. *MIC60* KO cells show a drastic reduction of protein levels of all MICOS components. **C** Quantification of CJs per mitochondrion in WT, *MIC10* and *MIC60* KO HAP1 cells using EM represented as boxplot (2 independent experiments, 24-40 mitochondria for each experiment). \*\*\*\**P* < 0.0001 (using unpaired Student’s *t*-test). **D** Quantification of cristae per mitochondrion in WT, *MIC10* and *MIC60* KO HAP1 cells using EM represented as boxplot (2 independent experiments, 24-40 mitochondria for each experiment). \*\*\*\**P* < 0.0001 (using unpaired Student’s *t*-test). **E** Comparison of oxygen consumption rates of basal, proton leak, maximum and non-mitochondrial respiration in WT and *MIC10* KO cells obtained from 3 independent experiments. \**P* = 0.034 for basal (using one sample *t-*test) and \**P* = 0.018 for maximum respiration (using unpaired Student’s *t*-test). **F** Comparison of oxygen consumption rates of basal, proton leak, maximum and non-mitochondrial respiration in WT and *MIC60* KO cells obtained from 3 independent experiments. \**P* = 0.01 for basal (using one sample *t-*test), \*\*\**P* = 0.0004 for maximum respiration and \**P* = 0.012 for non-mitochondrial respiration (using unpaired Student’s *t*-test).

### CJs are dispensable for regular arrangement of MIC60 in the IBM

MICOS is a large oligomeric complex present at CJs. Using STED (stimulated emission depletion) super-resolution images of fixed WT cells, we show a regularly spaced symmetrical arrangement of MIC10 and MIC60 punctae along the IBM (Fig 2A) consistent with earlier reports (Jans et al, 2013; Stoldt et al, 2019). We determined the distance between the two consecutive puncta of MIC10 and MIC60 in a mitochondrion and called it interpunctae distance (IPD). The IPD between two consecutive punctae was in the range of 280 nm both for MIC10 and MIC60 (Fig 2B) confirming that both subunits assemble into the MICOS complex in WT cells. Since *MIC60* deletion lead to reduction of all proteins of MICOS, while loss of *MIC10* still preserves MIC60 (Fig 1B), we asked whether MIC10 is required for the punctae-like appearance of MIC60. Deletion of MIC10, albeit leading to a loss of CJs, did not disturb the regular arrangement of MIC60 punctae along the mitochondrial length (Fig 2A). In line with this, the IPD of MIC60 in control and *MIC10 KO* cells was not significantly different (Fig 2B). Hence, lack of CJs and loss of the MICOS subcomplex of MIC10/13/26/27 did not alter the symmetrical arrangement of MIC60, concluding that CJs are not necessary for regular spacing of MIC60 in the IBM (Fig 2C). This supports the conclusion that MIC60 is positioned at uniform distances in IBM representing a docking and scaffolding platform for other MICOS subunits (Fig 2C) such as MIC10 for CJs formation, consistent with an earlier report in bakeŕs yeast (Friedman et al, 2015).

**Figure 2.**
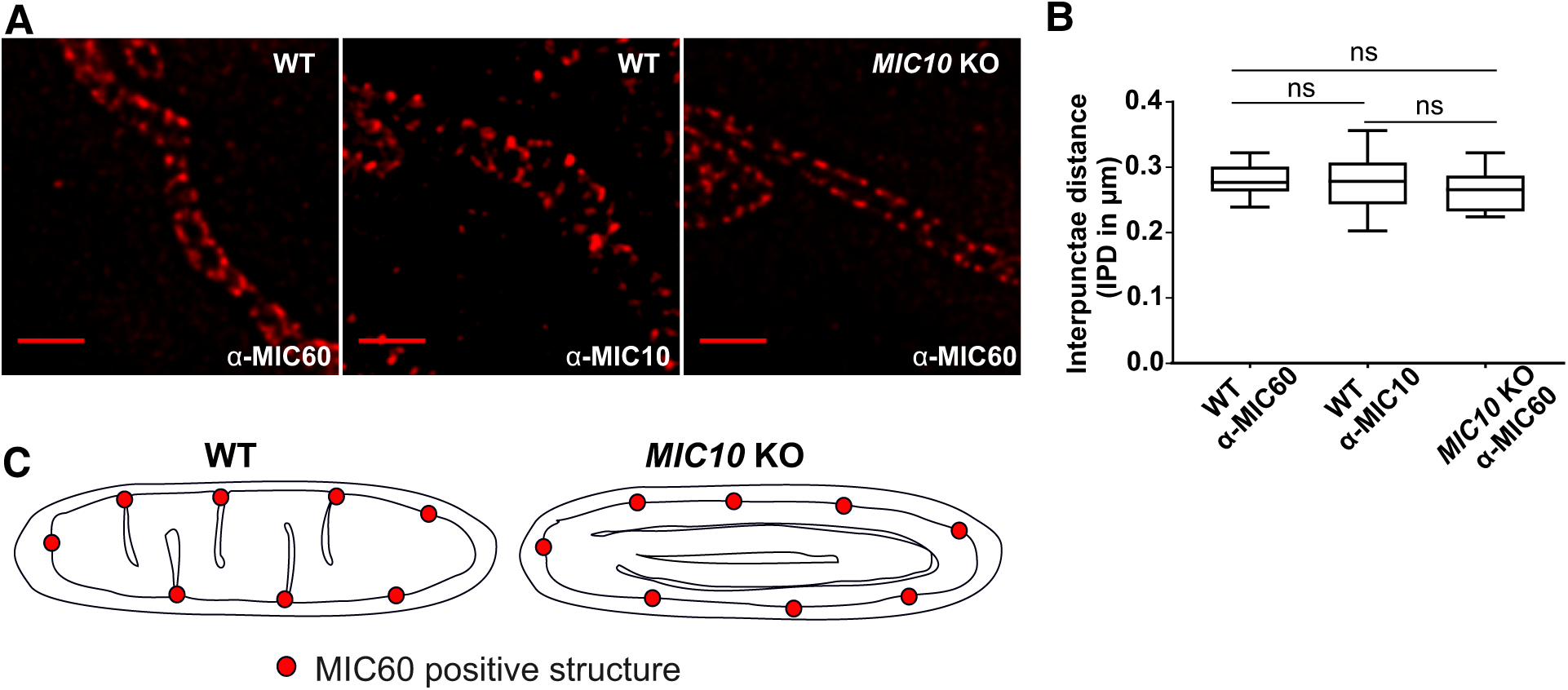
MIC60 assembled as regularly spaced punctae in absence of CJs. **A** Representative STED super-resolution images of WT HAP1 or *MIC10* KO HAP1 cells immunostained with indicated antibodies (MIC60 or MIC10). Scale bar 1 µm. **B** Quantification of mitochondrial interpunctae distances (IPD) between MIC60 or MIC10 punctae (2 independent experiments, 5-10 mitochondria for each experiment). ns, not significant, unpaired Student’s *t*-test. **C** Scheme representing MIC60 marking the nascent sites of CJs formation independent of MIC10.

### Crista junction proteins show a markedly reduced mobility in the inner membrane and loss of *MIC13* affects the mobility of distinct inner membrane proteins

We asked how mobile the two core MICOS subunits MIC60 and MIC10 are compared to other membrane proteins localized to different mitochondrial subcompartments. We constructed GFP-tagged versions of MIC60/MIC10, TOMM20, TIMM23 and ATP5I, established markers of CJs, OM, IBM and CM respectively (Fig 3A; EV4), and performed Fluorescence-Recovery After-Photobleaching (FRAP) experiments. For TOMM20, TIMM23 and ATP5I we observed substantially shorter T^1/2^ recovery times and higher mobile fractions than for MIC10 and MIC60, demonstrating that CJ proteins are much more restricted in movement compared to membrane proteins of other mitochondrial complexes present in various subcompartments (Fig 3B, 3D, EV1). In line with this, MIC60 was reported to show restricted diffusion in the inner membrane compared to outer membrane proteins in another study (Appelhans & Busch, 2017). To know whether the mobility of these proteins depends on the presence of a fully assembled MICOS complex, which is essential for formation of CJs, we generated *MIC13* KO Hela cells, as Hela cells are well suited for microscopy of mitochondria. Consistent with prior reports (Anand et al, 2016; Guarani et al, 2015a), we observed a loss of MIC10, MIC26, MIC27, altered cristae morphology, and loss of CJs in *MIC13* KO Hela cells (Fig EV2). Upon deletion of *MIC13,* in particular the mobility of TIMM23 and MIC10 was altered but not of MIC60 (Fig 3C, 3D, EV1B, EV1D, EV1F, EV1G). The latter is consistent with our finding that MIC60 can arrange in regularly spaced punctae in the absence of other MICOS subunits. Interestingly, the loss of MIC13 decreases the mobile fraction of TIMM23 considerably. As in baker’s yeast Tim23 was shown to dynamically redistribute between the CM and the IBM in a manner dependent on mitochondrial protein import (Vogel et al, 2006), we propose that the decreased TIMM23 mobile fraction in cells lacking CJs is due to trapping of a fraction of TIMM23 in the CM providing experimental support for the role of CJs as ‘gates’ between the CM and the IBM and acting as diffusion barriers. Moreover, the mobility of MIC10 increases drastically in *MIC13* KO Hela cells compared to WT cells, in line with the view that MIC13 is required for specifically stabilizing MIC10 at the MICOS complex (Anand et al, 2016; Guarani et al, 2015a).

**Figure 3.**
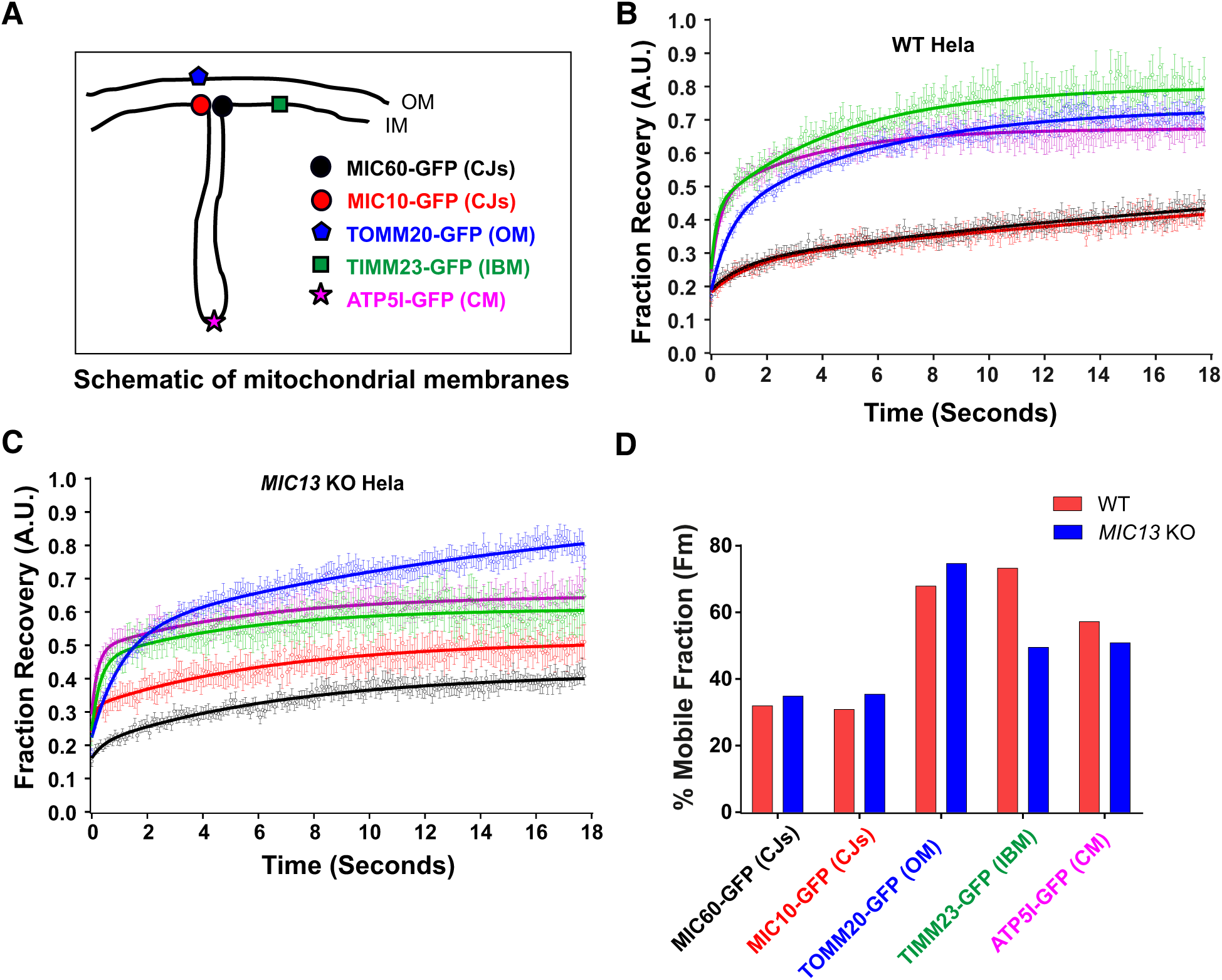
Mobility of crista junction proteins is restricted compared to proteins of other mitochondrial subcompartments and loss of MIC13 affects the mobility of distinct inner membrane proteins. **A** Scheme of investigated marker proteins, subjected to FRAP, located at different mitochondrial subcompartments. **B, C** FRAP curves (curve fitted) of WT **(B)** and *MIC13* KO Hela cells **(C)** expressing GFP-tagged fusion proteins of MIC60 (CJs), MIC10 (CJs), TOMM20 (OM), TIMM23 (IBM) and ATP5I (CM). Average FRAP curves (intensities in arbitrary units) for each marker were obtained (3 independent experiments, 6-10 mitochondria for each experiment). Error bars for each time point representing SEM are shown. **D** Histogram showing average percentage of mobile fractions, obtained from curve fits of all mitochondria expressing of GFP-tagged versions of MIC60 (CJs), MIC10 (CJs), TOMM20 (OM), TIMM23 (IBM) and ATP5I (CM) in WT and *MIC13* KO Hela cells.

To study the mobility of MIC10 and MIC60 by a different approach we used single particle tracking (Fig EV3). For this, we constructed the SNAP-tagged versions of MIC60 and MIC10 and confirmed their functionality as revealed by coimmunoprecipitation using anti-MIC13 antibodies (Fig EV4AB), BN-PAGE analysis (Fig EV4D), or restoration of MIC13 levels upon MIC10 expression (Fig EV4C). This confirmed that the tagging of MIC10 and MIC60 with SNAP or GFP does not interfere with their function and proper incorporation into MICOS complex. Using single particle tracking data, we observed that on average MIC60-SNAP showed a smaller instantaneous diffusion coefficient (insD) compared to MIC10-SNAP (Fig EV3E), consistent with the view that MIC60 represents regularly arranged nascent sites priming CJ formation which has to be steadily positioned. Deletion of *MIC13* led to a significant shift of tracks towards long-range movements (increased instantaneous diffusion coefficient; insD) of both MIC10 and MIC60 suggesting an increased mobility in both cases (Fig EV3E). This is fully in line with the FRAP data and the interpretation that the MICOS complex is not fully assembled in *MIC13 KO* cells. Further, in order to obtain a better understanding of the dynamic behaviour of individual single particles, each track was further subdivided into subtracks based on confinement in increasing order of directionality as confined diffusion, subdiffusion, normal diffusion, or directed motion (Fig EV3F-N). The percentage of subtracks showing directed motion is significantly higher for MIC10 than for MIC60 in WT Hela cells suggesting a considerably higher directionality for MIC10 (Fig EV3FG). Additionally, directed motion of both MIC10 & MIC60 is significantly higher in *MIC13* KO cells compared to WT (Fig EV3FG), indicating that both subunits, when present in the fully assembled MICOS complex have lower directed motion compared to the situation when the MICOS complex exists in separate subcomplexes. Intriguingly, MIC10 directed motion in *MIC60* KOs is significantly reduced compared to control cells indicating that the presence of MIC60 is specifically required for the enhanced directed motion of MIC10 (Fig EV3F). This suggests that MIC10 is recruited to the MICOS complex via MIC60. We propose that MIC60 acts as a docking platform pioneering CJ formation, consistent with earlier findings in baker’s yeast showing that MIC60-GFP assembles into discrete spots in a strain lacking all MICOS subunits (Friedman et al, 2015).

### Crista junctions move dynamically within seconds

FRAP and SPT data provided insights about the mobility of MIC10 and MIC60 as a group of molecules or individual MIC10 and MIC60 molecules within the IBM, respectively. As we observe regularly arranged CJ punctae using immunostaining of MIC10 and MIC60 and STED nanoscopy (Fig 2A), we next asked how and at what timescale do individual CJs move. For this, we devised a novel method to perform live-cell STED super-resolution nanoscopy using MIC60-SNAP and MIC10-SNAP as markers of CJs. SNAP-tag, a well characterized protein tag binds covalently to silicone-rhodamine (Gautier et al, 2008; Keppler et al, 2003) and allows high-resolution imaging with minimal fluorophore bleaching (Lukinavicius et al, 2013). The functionality and the proper incorporation of MIC10- and MIC60-SNAP into the MICOS complex was verified as described above (Fig EV4). The MIC10-SNAP and MIC60-SNAP fusion proteins were arranged in the same regularly spaced punctae-like pattern across the IBM (Fig 4A, 4C) as shown for the endogenous proteins using immunostaining (Fig 2A), providing further evidence for proper localization of the tagged constructs. We were able to image MIC10- and MIC60-SNAP for a total of ∼15s at least every 2.6 seconds. While mitochondria as a whole stay relatively static during this observation time, we observed rapid movements of MIC10/MIC60-SNAP punctae during this short time period **(Movie 1, Movie 2)**, demonstrating that CJs show an unexpectedly pronounced dynamic behaviour at a timescale of seconds. We repeatedly visualized these movies to observe any regular pattern in movement of MIC10 or MIC60 within a mitochondrion. Firstly, we focused on MIC10-SNAP and manually tracked the individual punctae of MIC10 in the subsequent frames (Fig 4A, **Movie 3).** We observed numerous instances where two MIC10-SNAP punctae rapidly joined each other and split and re-joined at a timescale of seconds (Fig 4B, **Movie 3**). We quantified the occurrence of joining/splitting events per unit length of mitochondria and found that the number of joining and splitting events were similar in a given time, indicating that these events are balanced and possibly coupled (Fig 5F). MIC60-SNAP showed a very similar pattern of joining and splitting (Fig 4D, 5E, **Movie 4**) that occurred at similar temporal frequency as MIC10-SNAP. We conclude that fully assembled MICOS in WT cells representing CJs undergo continuous cycles of joining and splitting in a rapid and balanced manner.

**Figure 4.**
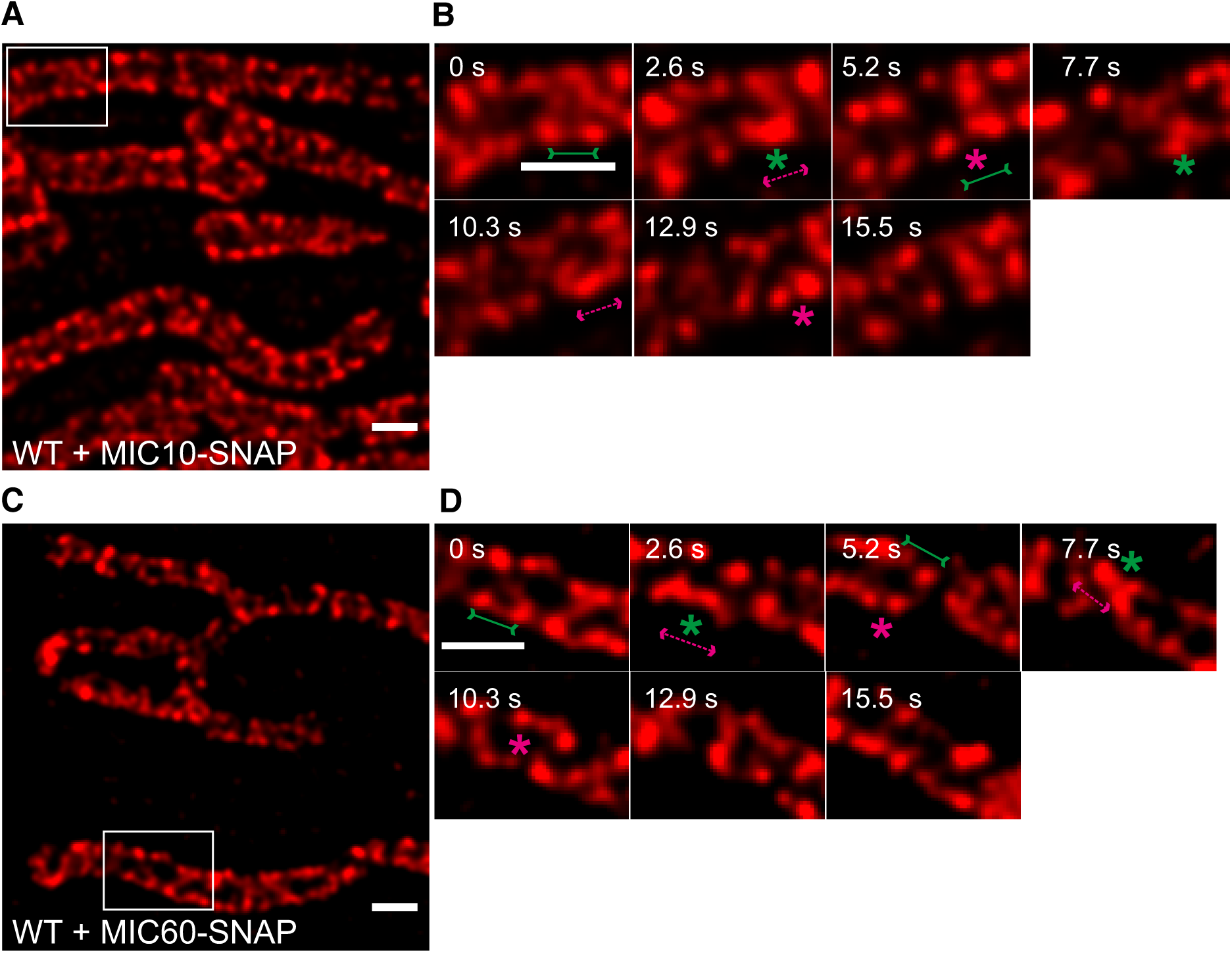
Crista junctions marked by MIC60-SNAP and MIC10-SNAP dynamically join and split within a mitochondrion. **A, C** Representative live-cell STED super-resolution image (t=0 s) showing WT Hela cells expressing MIC10-SNAP **(A)** and MIC60-SNAP **(C)** stained with silicone-rhodamine. Box in **A** and **C** mark selection shown as a zoom in panel **B** and **D** respectively. Scale bar 0.5 µm. **B, D** Time-lapse image series of a mitochondrion expressing MIC10-SNAP **(B)** or MIC60-SNAP **(D)** (2.6 s/frame). Green and magenta asterisks show joining and splitting events of MIC10-SNAP and MIC60-SNAP, respectively. Green arrows pointing inward connected by solid line and magenta arrows pointing outward connected by dotted line show sites of imminent joining and splitting events, respectively. Scale bar 0.5 µm.

**Figure 5.**
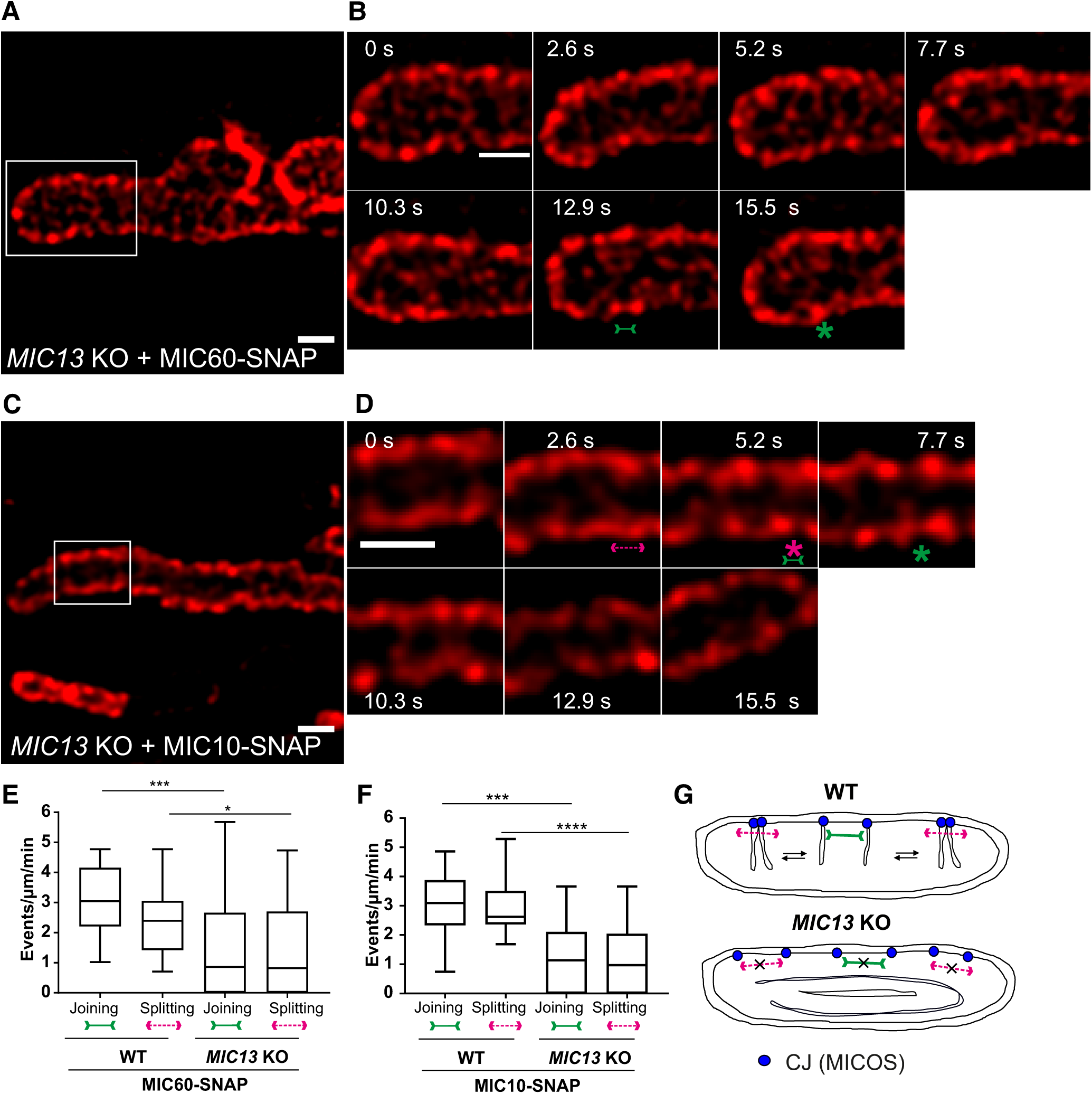
Joining and splitting of crista junctions occur in a MIC13-dependent manner. **A, C** Representative live-cell STED super-resolution image (t=0 s) showing *MIC13* KO Hela cells expressing MIC10-SNAP **(A)** or MIC60-SNAP **(C)** stained with silicone-rhodamine. Box in **A** and **C** mark selection shown as a zoom in panel **B** and **D** respectively. Scale bar 0.5 µm. **B, D** Time-lapse image series of a mitochondrion expressing MIC10-SNAP **(B)** and MIC60-SNAP in *MIC13* KO cells **(D)** (2.6 s/frame). Green and magenta asterisks show joining and splitting events of MIC10-SNAP and MIC60-SNAP, respectively. Green arrows pointing inward connected by solid line and magenta arrows pointing outward connected by dotted line show sites of imminent joining and splitting events, respectively. Scale bar 0.5 µm. **E** Quantification of joining and splitting events of CJs in WT Hela and *MIC13* KO cells expressing MIC60-SNAP (3 independent experiments, 5-7 mitochondria for each experiment). \*\*\**P* = 0.0004 (joining events WT vs. *MIC13* KO) and \**P* =0.019 (splitting events WT vs. *MIC13* KO), unpaired Student’s *t*-test. **F** Quantification of joining and splitting events of CJs in WT Hela and *MIC13* KO cells expressing MIC10-SNAP (3 independent experiments, 5-8 mitochondria for each experiment). \*\*\**P* = 0.0002 (joining events WT vs. *MIC13* KO) and \*\*\*\**P* < 0.0001 (splitting events WT vs. *MIC13* KO), unpaired Student’s *t*-test. **G** Scheme representing the dynamic nature of CJs in WT and *MIC13* KOs cells.

### CJs undergo joining and splitting event in a MIC13-dependent manner

To test whether the dynamic behaviour of MIC60-SNAP punctae depends on the presence of a fully assembled MICOS complex, we analysed dynamics of MIC60-SNAP in *MIC13* KO cells. We again observed the regular arrangement of MIC60-SNAP punctae in the IBM of mitochondria in *MIC13* KO cells (Fig 5A) consistent with our observation that MIC60 forms regularly arranged punctae also in the absence of MIC10 (Fig 2). MIC60-SNAP punctae showed dynamic movements in *MIC13* KO cells (Fig 5B, **Movie 5**), however, this occurred at a markedly reduced frequency when compared to WT Hela cells (Fig 5B, 5E, **Movie 5**). This indicates that, although the regular arrangement of MIC60 in principle is unaltered, the movement of MIC60-positive punctae reflected normally by balanced joining and splitting events is drastically impaired in the absence of CJs in *MIC13 KO* cells. We also overexpressed MIC10-SNAP in *MIC13* KOs and found that MIC10-positive punctae are dynamic, yet that joining and splitting events were markedly reduced as well (Fig 5C, 5D, 5F). These observations indicate that the frequency of regular joining and splitting of CJs depends on a functional MIC13-containing MICOS complex, which would be needed for cristae to emerge from CJs (Fig 5G).

To further corroborate our findings on CJ dynamics, we decided to analyse the movement of MIC13-SNAP, another marker for CJs, in WT Hela cells. Different from MIC60 or MIC10, we observed that the staining pattern of MIC13-SNAP not only marked regularly arranged punctae in the IBM but also appeared to label transverse bridges across mitochondria resembling cristae membranes (Fig 6A). Apparently, MIC13-SNAP is dually localized to both CJs and cristae. We cannot rule out that the partial localization of MIC13 to cristae is due to excess of MIC13-SNAP in the IM. Nevertheless, in this context this would even beneficial as it can help us to visualize cristae and CJs at the same time. We analysed the movement of MIC13-SNAP in live-cell STED nanoscopy and observed rapid dynamics of both CJs and cristae membranes at a timescale of seconds **(Movie 6).** We not only observed joining and splitting events of CJs as shown by MIC10-SNAP and MIC60-SNAP, but also rapid movement of cristae attached to these CJs (Fig 6B, 6D). Upon careful analysis we observed several instances showing that, when two CJs join, they bring adjoining cristae close by to fuse along near the IBM (Fig 6B, from 0s to 2.5s, **Movie 7**). Moreover, we observe other occasions where cristae were apparently fusing along the length of other cristae, sometimes forming structures resembling the letters X and Y (Fig 6B, at 5 s and 14.9 s, respectively). In many instances, fusion events are coupled to a subsequent apparent fission event at or near the site of the prior fusion event (Fig 6B, arrow pointing outwards). To better visualize these events, we acquired images every 1.3 s and also found numerous examples of fusion and fission of cristae (Fig 6C, 6D, **Movie 8, Movie 9**). We propose that cristae undergo continuous remodelling in form of cycles of fusion and fission in a manner that is linked to CJ dynamics.

**Figure 6.**
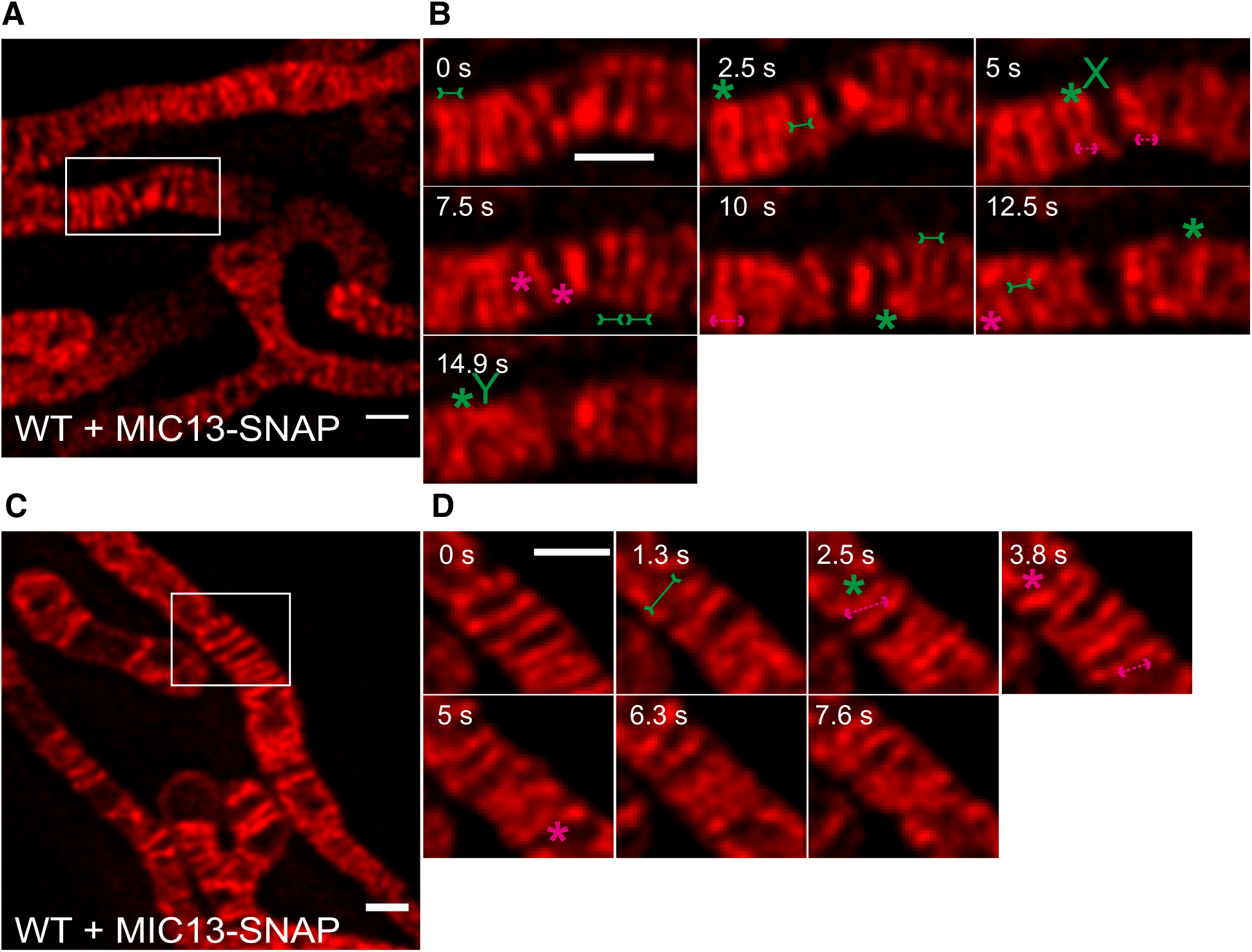
MIC13-SNAP shows that CJs and cristae undergo remodelling at a timescale of seconds. **A, C** Representative live-cell STED super-resolution image (t=0 s), showing WT Hela cells expressing MIC13-SNAP, from a time-series of images acquired at a time interval of 2.5 s **(A)** or 1.3 s **(C)** stained with silicone-rhodamine. Box in **A** and **C** mark selection shown as a zoom in panel **B** and **D** respectively. Scale bar 0.5 µm. **B, D** Time-lapse image series of a mitochondrion expressing MIC13-SNAP imaged at a time interval of 2.5 s/ frame **(B)** or 1.3 s/ frame **(D)**. Green and magenta asterisks show fusion and fission cycles of cristae marked by MIC13-SNAP. Green arrows pointing inward connected by solid line and magenta arrows pointing outward connected by dotted line show sites of imminent joining/fusion and splitting/fission events, respectively. A cristae fusion event that forms a structure resembling letter ‘X’ is marked in B at 5 s.

### Cristae undergo dynamic remodelling by continuous cycles of fusion and fission

To substantiate our observation of cristae fission and fusion using MIC13-SNAP, we obtained SNAP tagged versions of *bona fide* cristae markers, ATP5I and COX8A of F_1_F_O_ ATP synthase and complex IV, respectively. Hela cells expressing ATP5I-SNAP and COX8A-SNAP were imaged by live-cell STED super-resolution nanoscopy. The STED images of both ATP5I and COX8A showed transversely spanning bridges across the mitochondria (Fig 7A, 7C), consistent with electron micrographs of cristae in Hela cells. Cristae were rapidly changing their position in consecutive timeframes confirming that cristae are dynamic within mitochondria at a timescale of seconds (Fig 7B, 7D, **Movie 10**). Upon careful visualization, we observed several instances where cristae came close together and apparently fused either across the IBM (transverse type-fusion) or along their length (X- or Y-type fusion) (Fig 7B, 7D, **Movie 11, Movie 12**), consistent with our observations in MIC13-SNAP. Again, in most of the instances fusion events were accompanied by a subsequent fission event (Fig 7B, 7D). We quantified the rate of fusion or fission events across the unit length of mitochondria (µm) and determined that they occur in a balanced manner with ∼2 events per µm per min (Fig 7G). As opposed to the standard biochemistry textbook view, this makes it unlikely that cristae are constantly attached to CJs but rather can pinch-off transiently. Subsequently, they can re-fuse with the IBM or with other cristae or they can fuse across their length to make connections or even a branched network. Moreover, we checked whether the presence of a fully assembled MICOS is required for these continuous fusion and fission cycles using *MIC13* KO cells. We overexpressed COX8A-SNAP in *MIC13* KO cells and found that fusion and fission cycles are significantly reduced in cells lacking MIC13 (Fig 7E, 7G, **Movie 13**). Hence, a functional MICOS complex is required for these dynamic processes further supporting the view that CJ formation and dynamics are linked to cristae membrane dynamics.

**Figure 7.**
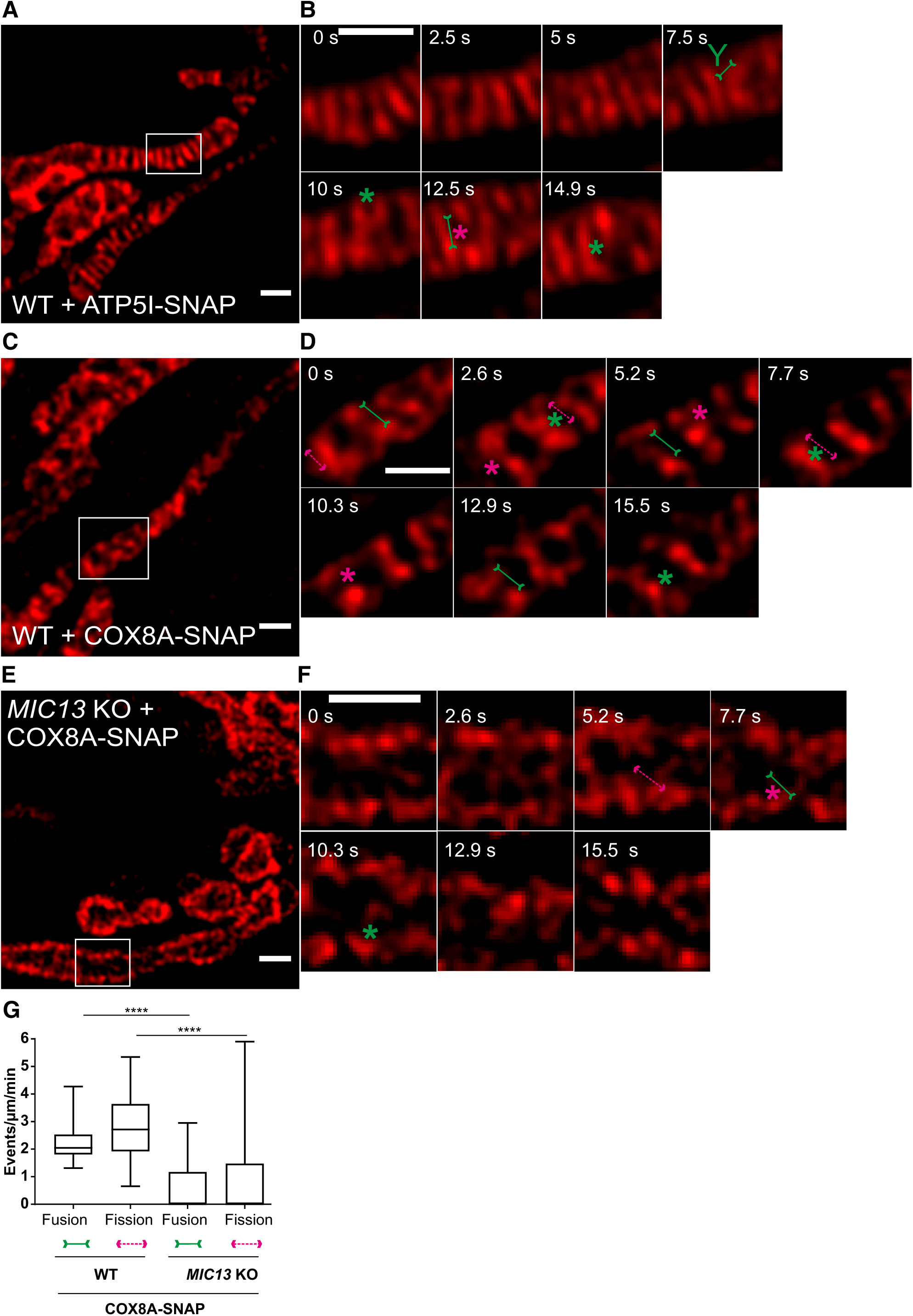
Cristae undergo balanced fusion and fission events in a MICOS-dependent manner. **A, C** Representative live-cell STED super-resolution image (t=0 s) showing WT Hela cells expressing ATP5I-SNAP **(A)** or COX8A-SNAP **(C)** stained with silicone-rhodamine. Box in **A** and **C** mark selection shown as a zoom in panel **B** and **D** respectively. Scale bar 0.5 µm. **B, D** Time-lapse image series of a mitochondrion expressing ATP5I-SNAP **(B)** or COX8A-SNAP **(D)** (∼2.5 s/frame). Green and magenta asterisks show fusion and fission events of cristae marked by ATP5I-SNAP or COX8A-SNAP, respectively. Green arrows pointing inward connected by solid line and magenta arrows pointing outward connected by dotted line show sites of imminent joining and splitting events, respectively. **E** Representative live-cell STED super-resolution image (t=0 s) showing *MIC13 KO* Hela cells expressing COX8A-SNAP stained with silicone-rhodamine. Box in **E** mark selection shown as a zoom in panel **F** respectively. Scale bar 0.5 µm. **F** Time-lapse image series of a mitochondrion expressing COX8A-SNAP in *MIC13 KO* **(D)** (2.6 s/frame). Rare fusion and fission events of cristae occurring in *MIC13* KO are marked by green and magenta asterisks respectively. Green arrows pointing inward connected by solid line and magenta arrows pointing outward connected by dotted line show sites of imminent fusion and fission respectively. **G** Quantification of cristae fission and fusion events in WT and *MIC13* KO Hela cells expressing COX8A-SNAP (3 independent experiments, 5-10 mitochondria for each experiment; \*\*\*\**P* < 0.0001, unpaired Student’s *t*-test).

### Cristae membrane dynamics allows redistribution of membrane potential indicating merging and content mixing of membranes

The timescales of intra-mitochondrial membrane dynamics, the balanced occurrence of these events, and the morphological features observed were very similar despite using a variety of protein markers (MIC10-, MIC60-, MIC13-, ATP5I- and COX8A-SNAP) for both CJs and cristae. This makes it unlikely that the observed cristae remodelling events are merely due to diffusion of different membrane protein complexes within a rather static IM. Still, we decided to use mitochondria-specific membrane dyes not labelling specific protein complexes. We further rationalized that cristae fusion events may result in an immediate change in the membrane potential ΔΨ of the two fusion mates, similar to the situation previously described for fusion between two mitochondria (Twig et al, 2008). Thus, we used TMRM, a dye labelling the inner membrane in a membrane-potential manner. Performing STED with TMRM, as opposed to SNAP-tagged proteins, is challenging due to its high sensitivity to photo-bleaching. Nevertheless, we could record movies for a similar time period as we did with SNAP-tagged protein markers using STED nanoscopy (Fig 8A, 8B). TMRM labelling showed transversely arranged cristae that were highly dynamic, consistent with our results for SNAP-tagged inner membrane proteins (Fig 8A, 8B). Moreover, we observed instances of immediate distribution of a TMRM signal from one crista (e.g. high-intensity signal at 1.5s in Fig. 8B) to another crista and IBM (lower-intensity signal at 3s), coinciding with the time of a cristae fusion event. Our observations strongly suggest that the location of the TMRM label within one lipid bilayer is rapidly altered in a short time period, implying a true membrane fusion event accompanied with content mixing. We also stained mitochondria with a non-potential specific dye, nonyl acridine orange (NAO), and employed another microscopy high-resolution technique, namely confocal laser scanning using a Zeiss Airyscan module. We observed that the cristae membrane was stained as transverse bridges in mitochondria and found instances of cristae dynamics at similar timescales (Fig 8C). We thus confirmed by two dyes and by a different imaging technique that cristae and CJs are highly dynamic and undergo continuous cycles of fusion and fission. Overall, we propose a model termed *<u>Cri</u>*stae-*<u>F</u>*ission-*<u>F</u>*usion’ (CriFF). In this unified model, we illustrate the observed fusion and fission cycles of cristae and how they are apparently linked to the observed dynamics of CJs (Fig 8D).

**Figure 8.**
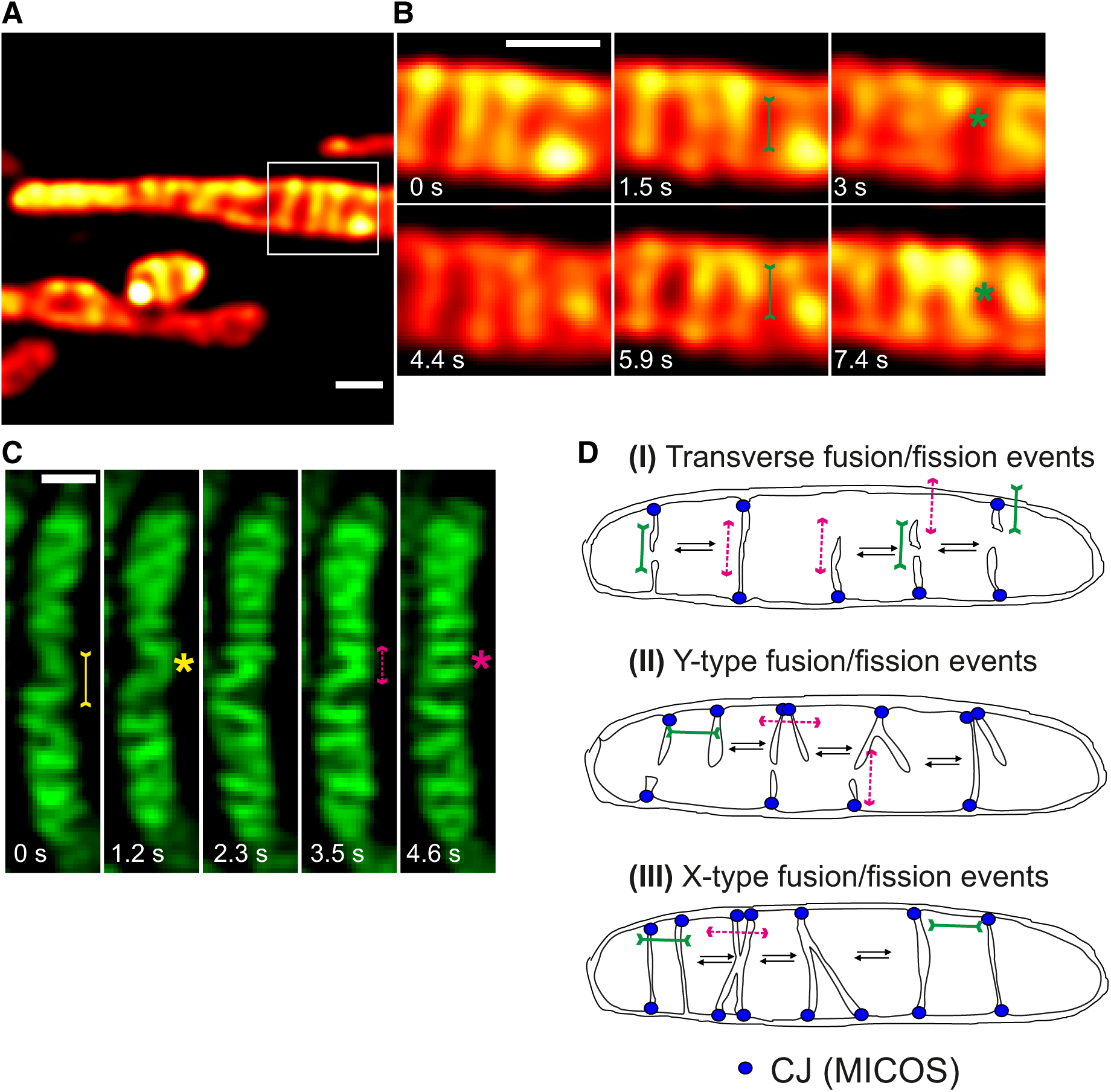
Cristae fission and fusion is corroborated using mitochondrial inner membrane-specific dyes. **A** Representative live-cell STED super-resolution image (t=0 s) showing WT Hela cells stained with TMRM. Box in **A** marks selection shown as a zoom in panel **B**. Scale bar 0.5 µm. **B** Time-lapse image series of a mitochondrion stained with TMRM (1.5 s/frame). Green asterisks show fusion events and redistribution of TMRM intensity from a high-intensity cristae to a low-intensity cristae (two such instances are shown at 3 and 7.4 s). Green arrows pointing inward connected by solid line at 1.5 and 5.9 s indicate imminent fusion events **C** Representative time-lapse images of WT Hela cells stained with NAO, and imaged using Airyscan microscope, indicate cristae fusion and fission dynamics. Yellow and magenta asterisks show fusion and fission events of cristae respectively. Yellow arrows pointing inward connected by solid line and magenta arrows pointing outward connected by dotted line show sites of imminent fusion and fission events, respectively. Scale bar 0.5 µm. **D** Schematic illustration of the ‘Cristae fission and fusion’ (CriFF) model and its link to CJ dynamics.

## Discussion

The idea that cristae must be able to undergo remodelling under certain physiological conditions has been around for half a decade, yet studying the dynamic behaviour of cristae membrane in real time was technically challenging due to limitations in optical resolution of diffraction-limited conventional light microscopy and due to the fact that electron microscopy can only capture fixed samples. In addition, the majority of studies using super-resolution techniques to study mitochondria have used fixed samples (Jans et al, 2013; Kukat et al, 2015; Stoldt et al, 2019). STED super-resolution microscopy of living cells has been utilized so far to visualize other cellular organelles, such as ER or Golgi (Schroeder et al, 2018; Shin et al, 2018). To date, only one study illustrated the power of Hessian structured illumination microscopy by giving an example in a single mitochondrion indicating cristae dynamics (Huang et al, 2018). Here we provide several lines of evidence demonstrating for the first time that CJs and cristae undergo fission and fusion cycles in a MICOS-dependent manner. These findings are based on the application of different advanced imaging techniques such as live-cell STED, SPT, FRAP and confocal Airyscan microscopy, which were applied to a wide variety of protein markers to visualize distinct mitochondrial substructures (MIC60-SNAP, MIC10-SNAP, MIC13-SNAP, ATP5I-SNAP, COX8A-SNAP), as well as two dyes labelling the inner membrane. We further obtained three mammalian knockout cell lines lacking *MIC10*, *MIC60* and *MIC13*, respectively, and expressed the various constructs of SNAP- and GFP-tagged mitochondrial IM proteins in those and wild type cells to reveal the functional role of the MICOS complex in this regard. Our improved tagging and imaging conditions allowed us to achieve a time resolution of up to 1.2 - 2.6 s per frame revealing that CJs and cristae dynamics occurs at very similar timescales (seconds) and both depend on the MICOS-complex.

The idea of MIC60 as a central pioneer of the MICOS complex was proposed from experiments in baker’s yeast as reintroducing MIC60-GFP into a strain lacking all MICOS subunits, shows MIC60-GFP to be assembled into discrete spots (Friedman et al, 2015). This in addition to our results corroborates that MIC60 is the pioneering subunit of the MICOS complex positioning MICOS assembly and CJs formation. Overall, we propose the ‘*<u>Cri</u>*stae-*<u>F</u>*ission-*<u>F</u>*usion’ (CriFF) model which integrates our observations as follows (Fig 8D). (1) MIC60 serves as a docking site for the formation of CJs via recruitment of other subunits of the MICOS complex. (2) Once the MICOS complex is fully assembled, CJs are the sites for cristae to grow and emerge from the IBM. This does not exclude that the MICOS complex can dynamically change its composition or assembly-state. (3) CJs can dynamically join and split. (4) Cristae are mainly, but not necessarily, attached to CJs. (5) Cristae can undergo transient pinching-off from CJs and rejoining at the same CJ or another one (transverse or neighbouring CJ; Fig 8Di). The formation of separated cristae subcompartments is consistent with another study showing that, within the same mitochondrion, individual cristae can maintain a stable membrane potential over time and that neighbouring cristae can maintain disparate ΔΨ (preprint: Wolf et al, 2019). This is also consistent with our FRAP data showing that TIMM23-GFP shows a reduced mobility upon loss of CJs, presumably by enhanced trapping in the CM. (6) Formation of Y-type cristae fusion may be preceded by CJ joining and subsequent cristae membrane remodelling (Fig 8Dii). This would represent one possibility to link CJ dynamics and cristae remodelling. (7) X-type cristae fusion could represent preceding CJ splitting from a Y-type like cristae morphology or cristae fusion between cristae distant from CJs (Fig 8Diii). This clearly is a working model and will require further testing in future studies.

What could be the physiological advantage for cristae to undergo fusion and fission and why could impairment thereof lead to severe human diseases including mitochondrial encephalopathy with liver dysfunction as reported for mutations in *MIC13* (Godiker et al, 2018; Guarani et al, 2016; Russell et al, 2019; Zeharia et al, 2016)? Although future experiments will have to provide the answer to these questions, from the results shown here and in other studies, it is very clear that OXPHOS is impaired in cells lacking MICOS subunits such as MIC10, MIC60, MIC26, MIC27, or MIC13 (Anand et al, 2016; Guarani et al, 2015b; Koob et al, 2015; Weber et al, 2013). How can we explain this and what could comprise possible functions of highly dynamic cristae membranes? To answer this, it is worth to consider that cristae fission would result in transient cristae subcompartments, which are physically isolated from the rest of the inner membrane. This goes along with a reduced surface of the inner membrane that is directly accessible to the cytosol via passing the intermembrane space and the outer membrane. On the other hand, cristae fusion reverts this and leads to an increase in accessible IM surface. The former situation is expected to favour trapping of metabolites and protons within the intracristal space, whereas the latter would increase the accessible surface, which is expected to favour metabolite exchange. As both processes, decreasing vs. increasing IM surface, are critical for mitochondrial function we propose that cristae dynamics is a way to adapt between these opposing necessities. Moreover, cristae fission and fusion may help to distribute and mix proteins, lipids, and important metabolites in the mitochondrion. In summary, we propose that cristae fission and fusion (CriFF) has major implications for the following processes: (1) Proton trapping; fission of cristae at CJs, resulting in transient intramitochondrial vesicles, could prevent proton leakage from the intercristal space to the cytosol, thereby promoting ATP synthesis. This does not exclude other ways of proton trapping proposed earlier, including proton buffering via cardiolipin (Kates et al, 1993) and preferential electrostatic local concentration at highly curved cristae tips (Strauss et al, 2008). Interestingly, pH differences within the intermembrane space have been reported, as the pH near ATP5I-SEcGFP (Superecliptic GFP, pH sensitive version of GFP), residing at cristae tips, was shown to be 0.3 higher compared to COX8A-SEcGFP (Rieger et al, 2014) located in the cristae. This was proposed to reflect a concentration gradient of protons, which dilutes at the cristae tips, which appear to act as a sink. (2) Content mixing; cristae fission and fusion may promote metabolite exchange including ADP, which was proposed to be limiting under certain conditions for OXPHOS (Mannella et al, 2013). (3) Intra-mitochondrial quality control by protein content mixing; ongoing fusion and fission dynamics of cristae may serve to mix partially damaged protein complexes, such as ETC complexes, in the CM to allow intra-mitochondrial complementation and protein quality control by proteases. (4) Biogenesis of membrane protein complexes; continuous mixing of newly imported protein subunits (nuclear as well as mtDNA encoded) may promote efficient assembly of membrane protein complexes, in particular when considering that the cristae membrane is very densely packed with proteins. This view is consistent with hybrid cristae, which were reported to be formed upon mitochondrial fusion, following PEG fusion of cells expressing mitochondria with different tags to respiratory chain complexes (Wilkens et al, 2013). (5) Lipid remodeling, mixing, and exchange; cristae dynamics could promote exchange of lipids between organelles and OM and IM and optimize cardiolipin synthesis. Contact sites between the IM and the OM, defined by the MICOS complex, were shown to coordinate the synthesis of phosphotidylethanolamine by Psd1 to dictate lipid remodeling in mitochondrial membranes (Aaltonen et al, 2016). (6) Other processes likely to be affected by altered cristae dynamics could include redox homeostasis, thermogenesis, Ca^2+^ buffering, and iron sulfur biogenesis.

What are the molecular machineries required for these processes? We show that these processes are dependent on fully assembled MICOS complex in a MIC13-dependent manner. This and the fact that both, CJs and cristae dynamics, occur at the similar timescales in a balanced manner, strongly suggest that they are mechanistically linked. Indeed, the MIC13-SNAP live-cell STED movies allow the simulatenous visualization of CJs and cristae supporting this view. Still, we cannot fully exclude that the MICOS complex has two independent functions explaining our observation equally well. Further experiments will provide insights into the bioenergetic parameters and the molecular mechanisms of these fascinatingly fast and continuously occurring dynamics of CJs and cristae.

In conclusion, live-cell STED super-resolution nanoscopy in combination with biochemical and genetic methods has expanded our view on the dynamics of IM remodelling at a nanoscopic level in real-time. Future research will provide further insights into the regulation, physiology, and pathophysiology of these newly revealed dynamic processes occurring within mitochondria.

## Materials and Methods

### Cell culture, transfection and generation of knockout cell lines

Hela and HEK293 cells were maintained in DMEM (Sigma-Aldrich) supplemented with 10 % fetal bovine serum (PAN Biotech), 2 mM glutaMAX (Gibco), 1 mM sodium pyruvate (Gibco) and penstrep (Sigma-Aldrich, penicillin 100 units/ml and streptomycin 100 µg/ml) whereas HAP1 cells were cultured using Isocovés modified DMEM media (IMDM) supplemented with 20 % fetal bovine serum (PAN Biotech), 2 mM glutaMAX (Gibco) and penstrep (Sigma-Aldrich, penicillin 100 units/ml and streptomycin 100 µg/ml). Cells were grown in incubator with 37°C and 5% CO_2_. All cell lines were tested negative for possible mycoplasma contamination. Hela and HAP1 cells were transfected with 1 µg of corresponding plasmid using GeneJuice® (Novagen) according to the manufacturer’s instructions. In case of SNAP-tag constructs, 0.2 µg of MitoGFP (matrix-targeted) and 1 µg of corresponding SNAP-tag were co-transfected. HEK293 cells were grown on a large-scale in 10 cm dishes and transfected with 10 µg of respective plasmids were used for biochemical experiments. *MIC13* KO Hela cells were generated using CRISPR/Cas method as described before (Anand et al, 2016). *MIC10* and *MIC60* KO HAP1 cells along with WT cells were custom-made by Horizon (UK).

### Molecular cloning

Human MIC60, MIC10, MIC13, ATP5I and TOMM20 were cloned into pSNAPf vector (NEB) using Gibson assembly cloning kit (NEB). COX8A-SNAP vector was commercially obtained from NEB. MIC60 and MIC10 were cloned into pEGFPN1 using restriction digestion by *Xho*I and *Bam*H1 followed by ligation. Human TOMM20, TIMM23, ATP5I were cloned into pEGFPN1 vector using Gibson assembly cloning kit (NEB) to acquire their respective GFP-tagged versions.

### SDS electrophoresis and western blotting

For preparing samples for western blotting, corresponding cells were collected and proteins were extracted using RIPA lysis buffer. The amount of solubilized proteins in all the samples were determined using the Lowry method (Biorad). 15% SDS-PAGE was performed, separated proteins were subsequently blotted onto a PVDF membrane, and probed with indicated antibodies: MIC10 from Abcam (84969), MIC13 (custom-made by Pineda (Berlin) against human MIC13 peptide CKAREYSKEGWEYVKARTK), MIC19 (Proteintech, 25625-1-AP), MIC25 (Proteintech, 20639-1-AP), MIC26 (Thermo-Fisher, MA5-15493), MIC27 (Atlas Antibodies, HPA000612), MIC60 (custom-made, Pineda (Berlin) against human IMMT using the peptide CTDHPEIGEGKPTPALSEEAS), SNAP-tag (P9310S, NEB), and β-tubulin (abcam, ab6046). Goat anti-mouse IgG HRP conjugated (ab97023) and goat anti-rabbit IgG HRP conjugated antibody (Dianova, 111-035-144) were used as secondary antibodies. Chemiluminescence was captured using a VILBER LOURMAT Fusion SL (Peqlab).

### Coimmunoprecipitation

For coimmunoprecipitation, isolated mitochondria from HEK293 cells overexpressing either MIC10-SNAP, MIC10-GFP, MIC60-SNAP or MIC60-GFP were used. Mitochondrial isolation was done according as described before (Anand et al, 2016). The coimmunoprecipiation experiment was performed using the protocol described in (Barrera et al, 2016) with following modification. The beads were incubated with 4 µg of MIC13 antibody (custom-made by Pineda (Berlin) against human MIC13 peptide CKAREYSKEGWEYVKARTK). During lysis of the mitochondria a detergent/protein ratio of 2 g/g was used.

### Isolation of macromolecular complexes by blue native gels

Mitochondria from HEK293 cells overexpressing MIC60-SNAP or MIC60-GFP were isolated as previously shown by (Anand et al, 2016). The experiment was performed as shown by (Anand et al, 2016) with the use of a detergent/protein ration of 2 g/g during solubilization.

#### Electron microscopy

HAP1 WT, *MIC10* KO and *MIC60* KO cells were grown on petri dishes, cells were washed with PBS, and fixed using 3% glutaraldehyde in 0.1M sodium cacodylate buffer, pH 7.2. After fixation, cells were collected in a small tube using a cell scraper and pelleted. These cell pellets were washed with 0.1M sodium cacodylate, pH 7.2 and subsequently embedded in 2% agarose. The pellets were stained using 1% osmium tetroxide for 50 min and 1% uranyl acetate/ 1% phosphotungstic acid for 1 h. The samples were dehydrated using graded acetone series and embedded in spur epoxy resin for polymerization at 65°C for 24 h. The ultrathin sections were prepared using microtome and the images were acquired using transmission electron microscope (Hitachi, H600) at 75V equipped with Bioscan model 792 camera (Gatan) and analysed with ImageJ software.

#### Cellular respiration measurements

All respiration measurements were performed using Seahorse XFe96 Analyzer (Agilent). The HAP1 cells were seeded in Seahorse XF96 cell culture plate (Agilent) at a density of 30,000 cells per well overnight. Next day, cells were washed and incubated in basic DMEM media (Sigma, D5030) supplemented with glucose, glutamine and pyruvate at 37°C in non-CO_2_ incubator 1 h prior to the assay. Mitochondrial respiration function was measured using Seahorse XF Cell Mito Stress Test kit (Agilent) according to the manufacturer’s instructions. Briefly, the delivery chambers of the sensor cartridge were loaded with oligomycin (F_1_F_O_-ATPase synthase inhibitor) or FCCP (uncoupler) or rotenone and antimycin (Complex I and Complex III inhibitors, respectively) to measure basal, proton leak, maximum and residual respiration. Cell number was normalized after the run using Hoechst staining. Data was analysed using wave software (Agilent).

#### Immunofluorescence staining

HAP1 cells were fixed with pre-warmed (37°C) 3.7 % paraformaldehyde for 15 min. After fixation, cells were washed 3 times with PBS, permeabilized with 0.15 % Triton-X 100 for 15 min, blocked using 10 % goat serum for 15 min followed by incubation with appropriate dilution of primary antibodies for 3 h at room temperature or overnight at 4°C. After washing thrice with PBS, samples were incubated at room temperature with appropriate secondary antibody for 1 h and washed three times with PBS before proceeding for microscopy. For STED super-resolution imaging primary antibodies used were against MIC60 (custom-made, Pineda (Berlin)), MIC10 (abcam, 84969). Goat anti-rabbit Abberior STAR 635P (abberior) was used as secondary antibody.

#### Quantification of mitochondrial interpunctae distance (IPD)

The longitudinal distance along the mitochondrial length between two successive MIC60 or MIC10 punctae is termed as interpunctae distance (IPD). The IPD between MIC60 and MIC10 punctae was calculated using the Image J software by manually drawing lines between two spots from the centre of the punctae and measured using the ‘Analyze’ function to calculate the length of that particular line. In order to avoid repetition of measuring the IPD, the length between the punctae was measured in a clockwise direction. If the edges of mitochondria containing the MIC60 or MIC10 spots were curved, a segmented line tool was used to measure the IPD. In rare cases where MIC60 spots were replaced by longitudinal bridges, the centre of the line was taken into consideration for defining the spot. The distance between MIC60 or MIC10 punctae per mitochondrion was calculated (from an average of 36-55 spots/mitochondrion) and a median interpunctae distance was obtained for that individual mitochondrion and plotted as boxplots.

#### 2D Live-STED (stimulated emission depletion) super-resolution nanoscopy

Cells transfected with corresponding SNAP-tags and stained with silicone-rhodamine dye (SNAP-cell 647-SiR (NEB) (3 µM)) were imaged at 37°C and 5 % CO_2_ mimicking cell culture incubator conditions in fluoroBrite DMEM media supplemented with 10 % fetal bovine serum (PAN Biotech), 2 mM glutaMAX (Gibco), 1 mM sodium pyruvate (Gibco) and penstrep (Sigma-Aldrich, penicillin 100 units/ml and streptomycin 100 µg/ml). For TMRM imaging, Hela cells were stained with 50 nM TMRM dye (Invitrogen) for 30 mins. Live-STED super-resolution nanoscopy was performed on Leica SP8 laser scanning confocal microscope fitted with a STED module. Before imaging the alignment of excitation and depletion laser was checked in reflection mode using colloidal 80 nm gold particles (BBI Solutions). Cells expressing the SNAP-tag and stained with SNAP-cell 647-SiR (NEB) were excited with a white light laser at an excitation wavelength of 633 nm. Images were collected using a hybrid detector (HyD) at an emission range from 640 nm to 730 nm using a 93X glycerol (N.A = 1.3) or 100X oil objective (N.A = 1.4) while using a pulsed STED depletion laser beam at 775 nm emission wavelength. The images were obtained at a zoom to acquire 9.7 × 9.7 µm area (12X magnification for 100X or 12.9 X magnification for 93X objective). Movies were obtained at a frame rate of ∼2.5 s/frame. Gating STED was used from 1 ns onwards in order to increase the specificity of the signal in experiments where silicone-rhodamine was used. Image processing post-image acquisition of live-cell SNAP movies was performed with Huygens Deconvolution software. Due to the strong depletion and the drastic decrease of emitting fluorophores in STED microscopy, the resulting images tend to have a lower signal to noise ratio. The increase of axial resolution, and by that reduction of out of focus blur, is not the only benefit of deconvolution. The other major advantages are increase in signal to noise ratio and resolution, which enhance image quality for 2D datasets as well, as shown before (Schoonderwoert et al, 2013).

#### Quantification of crista junction and cristae dynamics

Intramitochondrial joining/splitting events in case of MIC10- and MIC60-SNAP marking CJs and fusion/fission events in case of COX8A-SNAP marking cristae were quantified manually by observing the corresponding live-STED movies for a time span of ∼15 s obtained at a frame rate of ∼2.5 s/frame. The number of joining/splitting or fusion/fission events were divided by the mitochondrial length to yield corresponding number of events/ unit length (µm) of mitochondrion. Events/µm length/min of mitochondria was subsequently calculated. All quantifications were performed with 3 independent experiments using 5 to 10 mitochondria, which were all chosen from separate cells.

#### FRAP and associated quantification

FRAP experiments were performed on Leica SP8 using the FRAP module with Fly mode function switched on. Images were acquired with 40 X water objective (N.A = 1.1) using 25 X zoom. In order to avoid acquisition photobleaching, only 1-1.5% laser power of the Argon laser line at 488 nm was used to acquire the images using a PMT in green emission range. A square region of 0.7 × 0.7 µm was bleached using 100 % laser power at 488 nm. 10 pre-bleach images were acquired while 10 images were acquired during bleaching. 200 post-bleach images were acquired at a maximal possible frame rate of 88-89 ms/frame to monitor the recovery of fluorescence. After the images were acquired, quantification of the FRAP experiment was done in the following way: three different ROIs (Regions of Interest) were taken into consideration: 1) ROI^1^, an area where no mitochondria were found in the image, was used to perform background subtraction. 2) ROI^2^ was the area of mitochondria where the photobleaching was performed. 3) ROI^3^, another region of a separate mitochondria not subjected to FRAP, was used to obtain correction factor for acquisition photobeaching. ROI^2^(P) was the average of 10 pre-bleach measurements of ROI^2^ whereas ROI^3^(P) was the average of 10 pre-bleach measurements of ROI^3^. Hence, photobleach correction was performed by using the formula: ROI^2^-ROI^1^/ROI^3^-ROI^1^ and normalization was performed by using the formula: ROI^2^-ROI^1^/ROI^3^-ROI^1^ X ROI^3^(P)-ROI^1^/ROI^2^(P)-ROI^1^. All the mitochondria belonging to a particular condition from independent experiments were pooled and averaged for their FRAP curves. Standard error of mean (SEM) was plotted for all the pooled mitochondria for each condition.

Once the FRAP recovery values were obtained, they were fitted, using Graphpad Prism 7.04, by nonlinear regression two phase association model for all molecules. First and second phase association T^1/2^ recovery values were obtained for each condition when the curves were fitted by nonlinear regression two phase association model. Mobile fraction was calculated by using the formula: F_m_ = F_p_-F_0_/F_initial_-F_0_ where F_m_ denotes the mobile fraction, F_p_ denotes the fraction of fluorescence when the plateau is reached, F_initial_ is 1 and F_0_ is the fraction of initial fluorescence after the last pulse of photobleaching for all cases except MIC60-GFP, MIC10-GFP and TOMM20-GFP (in *MIC13* KO cells), where the Fp was calculated from the average of last 5 intensity values. Diffusion coefficients (D) were calculated using the formula D=0.25 X r^2^/T^1/2^, where r is the radius of the bleached ROI and T^1/2^ is the recovery time in seconds, as suggested before (Kang et al, 2012). We bleached a square ROI region of 0.49 µm^2^ in the mitochondria for our experiments. Hence, we used 0.49 µm^2^ as the area of the circle to calculate the radius.

#### Single particle tracking imaging and quantification

Cells transfected with the SNAP-tag and stained with low concentrations of SNAP-cell 647-SiR (NEB) (15 nM for MIC60-SNAP and 0.225 nM for MIC10-SNAP) were imaged in fluoroBrite DMEM media supplemented with 10 % fetal bovine serum (PAN Biotech), 2 mM glutaMAX (Gibco), 1 mM sodium pyruvate (Gibco) and penstrep (Sigma-Aldrich, penicillin 100 units/ml and streptomycin 100 µg/ml). Lower concentrations of silicone-rhodamine were used here compared to STED super-resolution imaging to allow for selective labelling of only few molecules in a mitochondrion. Movies were acquired on a Zeiss Elyra PS.1 microscope equipped with a 63x (NA = 1.46) objective lens in total internal reflection fluorescence (TIRF) mode, where highly inclined and laminated optical sheet (HILO) illumination was used, at a frame rate of 33 ms/frame for 1000 frames. The angle of the illuminating laser, EMCCD gain and laser intensity were manually adjusted to acquire the best signal to noise ratio (SNR). The first 50-100 frames of every movie were not used for analyses so that acquisition bleaching will additionally provide a sufficiently low concentration of single particles. For analyses of movies, a ROI was set around each cell and single particle tracking (SPT) was performed using the Fiji / ImageJ (Schindelin et al, 2012) plugin TrackMate (Tinevez et al, 2017) where the following TrackMate settings were used: Detector: Laplacian of Gaussian, Estimated blob diameter: 0.5 µm, Do sub-pixel localisation: Yes, Initial thresholding: No, Filter on spots: SNR above 0.4. For tracking spots, the LAP (Linear Assignment Problem) tracker with following settings worked well: Frame to frame linking: 0.3 µm, Gap-closing maximal distance: 0.5 µm, Gap-closing maximal frame gap: 5 frames. The filter on tracks function were used to only analyse tracks with more than 20 spots within a track. After analysis, the calculated mean square displacement (MSDs) and instantaneous diffusion coefficients (insD) of all tracks were loaded into the Fiji plugin “Trajectory Classifier” (Wagner et al, 2017). MSD analysis allows to determine the mode of displacement of particles over time. For the calculation of the MSD for every track and time point, the MATLAB class @msdanalyzer was used (Tarantino et al, 2014). Additionally the insD were calculated using insD=mean MSD / 4*dt (Giannone et al, 2010) where mean MSD were calculated using the first 4 MSD points of every track. Comparison of our results of insD of well characterized GPI-anchored proteins in our hands matched well with literature validating our SPT method and its quantification (data not shown). Beside for the calculation of the MSD and insD, tracks were analysed using the Fiji plugin Trajectory classifier (Wagner et al, 2017). The plugin classifies tracks generated with TrackMate into confined diffusion, subdiffusion, normal diffusion and directed motion in increasing order of directionality. As most tracks were relatively short, following settings for the classification were used: Minimal track length: 20, Window size: 10, Minimal segment length: 10. SPT data was obtained for each condition in 2 independent experiments from 2-5 cells in each experiment.

### Airyscan microscopy

Hela cells were stained with 100nM nonyl acridine orange for 1 h and imaged using Plan-Apochromat 100X/1.46 Oil DIC M27 objective on the Zeiss LSM 880 with Airyscan. Raw.czi files were processed into deconvoluted images using the Zen software automatically.

#### Statistics

Statistical analysis for different experiments were done in Graphpad Prism 7.04 using unpaired Student’s *t*-test except for comparison in Seahorse experiments where differences in basal oxygen consumption were compared by one sample *t-*test. Mean ± SEM was used for different experiments and sample size was not predetermined using any statistical methods.

## Acknowledgements

We thank T. Portugall and A. Borchardt for technical assistance in constructing the plasmids used in this study and in electron microscopy experiments, respectively. STED, FRAP and SPT experiments were performed at the Centre for Advanced Imaging (CAi) facility at HHU, Düsseldorf. We are also grateful to Prof. Wilhelm Stahl, Prof. Peter Brenneisen and Dr. Marcel Zimmermann for helpful and stimulating discussions. This work was supported by Deutsche Forschungsgemeinschaft (DFG) grant RE 1575/2-1 (AKK and ASR), SFB 974 Project B09 (ASR), Research Committee of the Medical faculty of Heinrich Heine University Düsseldorf FoKo-37/2015 (AKK) and Foko-02/2015 (RA & ASR), SFB 1208 Project Z02 (SWP) and grant WE 5343/1-1 (SWP).

## Author Contributions

AKK, RA and ASR developed the underlying concept of the study, designed the experiments and wrote the manuscript with comments from all other authors. AKK performed all experiments of STED super-resolution imaging with help from SH and SWP. RA performed all the biochemical experiments, and electron microscopy. FRAP experiments were done by AKK with help from SWP. TZ and AKK performed the SPT experiments while TZ performed quantification of SPT data using Image J with a MATLAB interface. JU characterized the *MIC13* KO cells and performed co-IP and BN-PAGE experiments. DMW, MS performed all experiments using Zeiss Airyscan imaging and analysis. ML and OSS contributed with scientific and critical inputs to the manuscript.

## Figure Legends

**Figure EV1.**
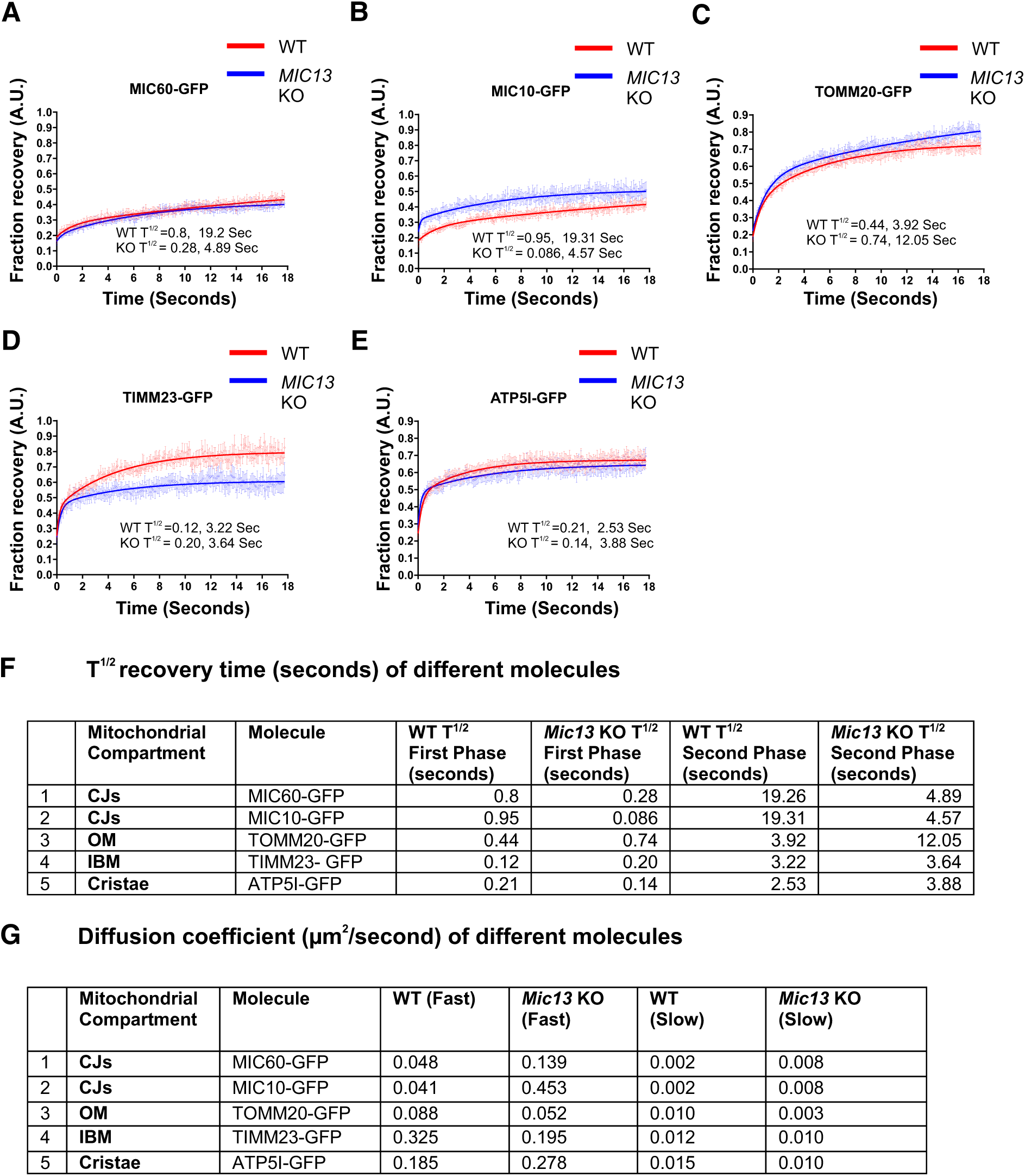
Detailed analysis of FRAP measurements of fusion proteins localizing to different mitochondrial subcompartments in WT and *MIC13* KO Hela cells. **A-E** FRAP curves (curve-fitted) of **A,** MIC60-GFP **B,** MIC10-GFP **C,** TOMM20-GFP **D,** TIMM23-GFP **E,** ATP5I-GFP in WT and *MIC13* KO Hela cells. Average FRAP curves (intensities in arbitrary units) for each protein were obtained (3 independent experiments, 6-10 mitochondria for each experiment). Error bars for each time point representing SEM are shown. **F, G** T^1/2^ recovery time **(F)** and diffusion coefficients **(G)** of indicated proteins.

**Figure EV2.**
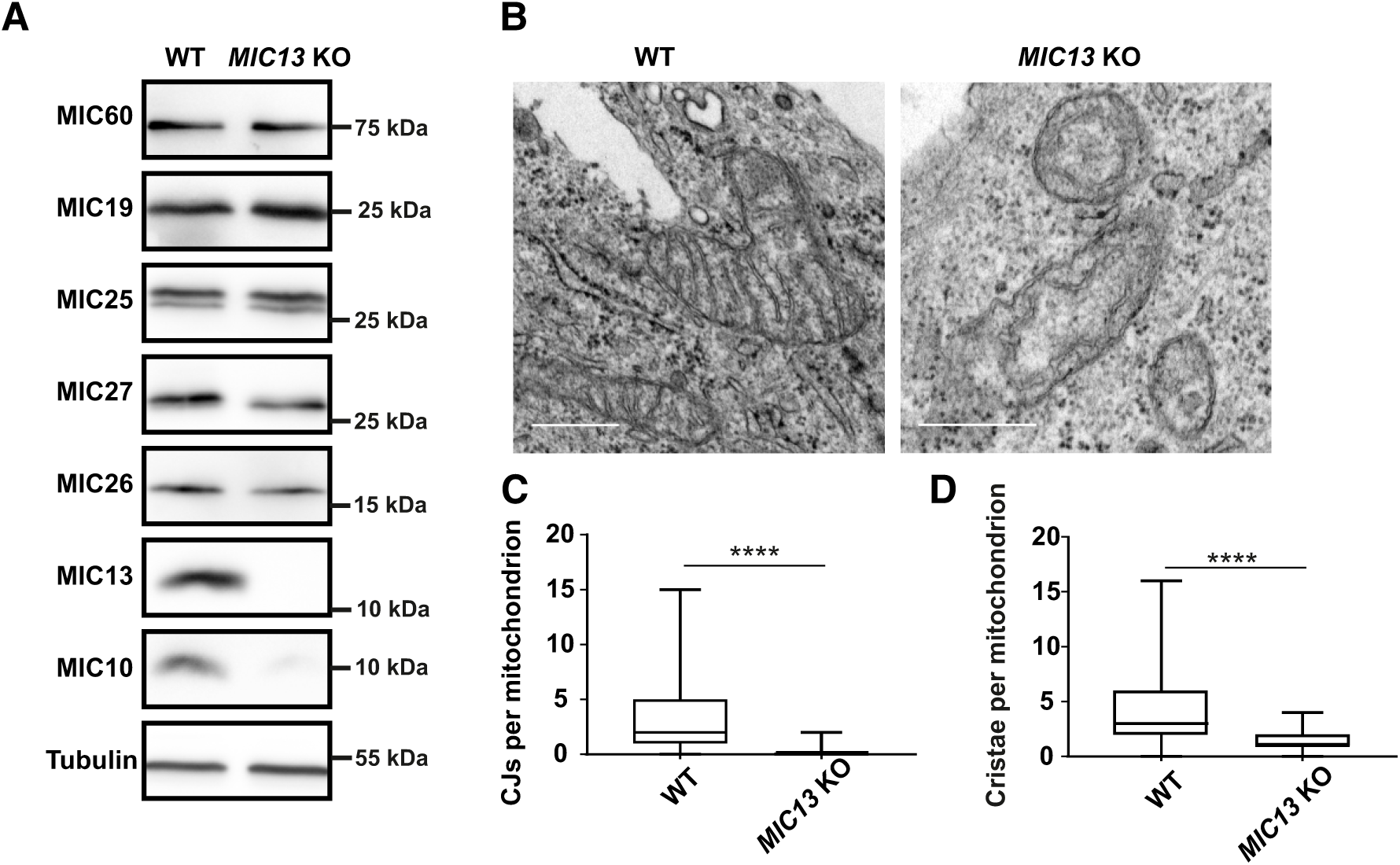
Deletion of *MIC13* in Hela cells leads to a loss of crista junctions. **A** Western blot analysis of WT and *MIC13* KO Hela cells showing a reduction of MIC10, MIC26 and MIC27 protein levels in *MIC13* KOs. **B** Representative electron micrographs of WT and *MIC13* KO Hela cells show loss of CJs in *MIC13* KOs. Scale bar 0.5 µm. **C** Boxplot showing quantification of CJs per mitochondrion in WT and *MIC13* KO Hela cells (2 independent experiments, 24-49 mitochondria for each experiment). \*\*\*\**P* < 0.0001 (WT vs. *MIC13* KO), unpaired Student’s *t*-test. **D** Boxplot showing quantification of cristae per mitochondrion in WT and *MIC13* KO Hela cells (2 independent experiments, 24-49 mitochondria for each experiment). \*\*\*\**P* < 0.0001, unpaired Student’s *t*-test.

**Figure EV3.**
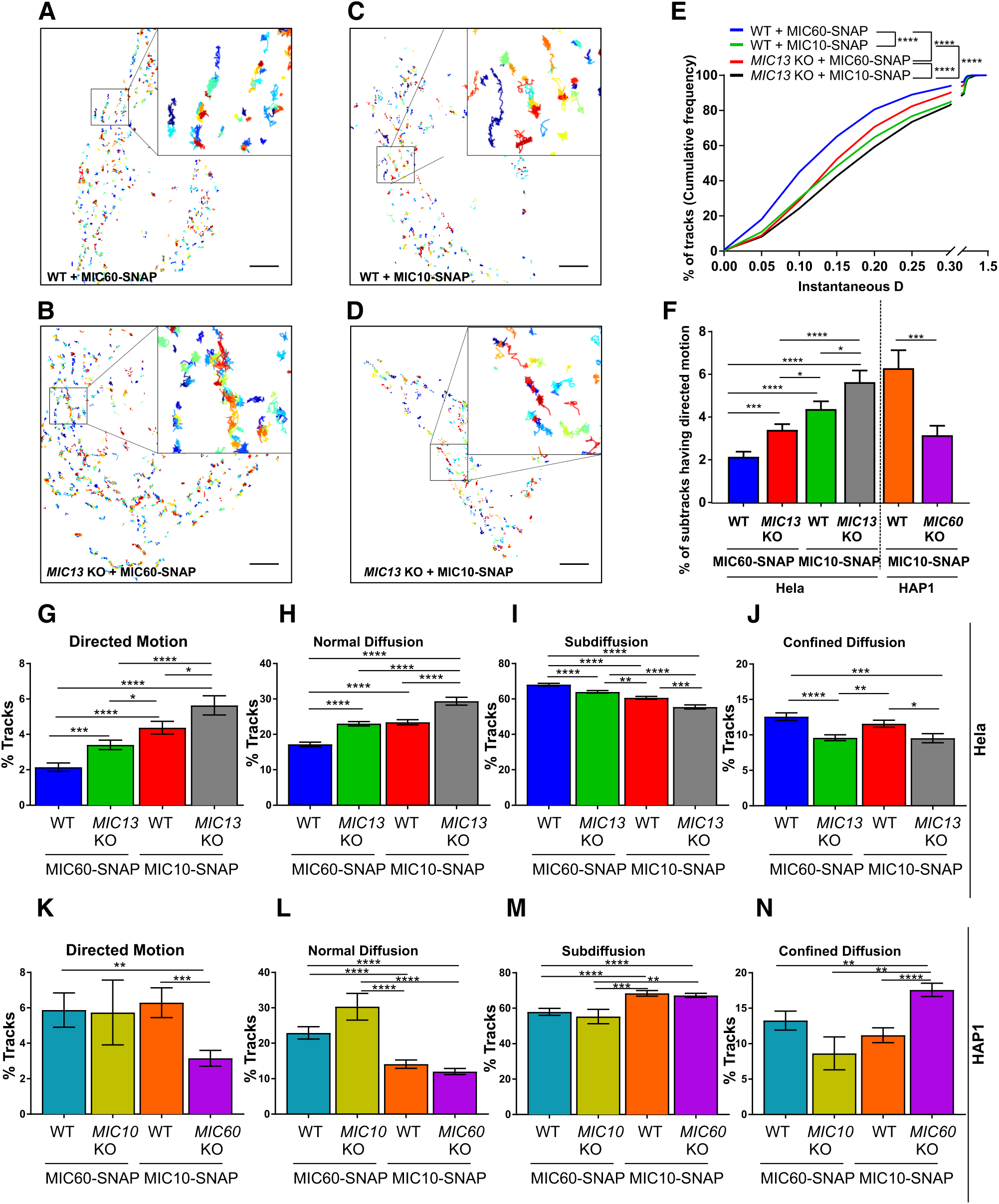
Single particle tracking reveals requirement of MIC60 for high directionality of MIC10. **A-D** Representative single particle tracks of cells expressing MIC60-**(A)** and MIC10-SNAP **(C)** in WT Hela cells and MIC60-**(B)** and MIC10-SNAP **(D)** in *MIC13* KO Hela cells, stained with silicone-rhodamine dye, and imaged at a rate of 33 ms/frame. Single tracks were color-coded according to temporal appearance. Insets in each image **(A-D)** show zoomed-in images of few representative tracks. **E** Cumulative frequency of tracks having corresponding values of instantaneous diffusion coefficients (insD) in WT and *MIC13* KO Hela cells expressing MIC60- and MIC10-SNAP stained with silicone-rhodamine. Number of tracks analysed: 2541, 2540, 3560 and 1441 in WT Hela cells expressing MIC60-SNAP and MIC10-SNAP, *MIC13* KO Hela cells expressing MIC60-SNAP and MIC10-SNAP, respectively. (\*\*\*\**P* ≤ 0.0001 for all possible comparisons, unpaired Student’s *t*-test). **F** Percentage of subtracks having directed motion in WT and *MIC13* KO Hela cells expressing MIC60- and MIC10-SNAP and WT and *MIC60* KO HAP1 cells expressing MIC10-SNAP. Number of corresponding subtracks analysed: 2561, 2443, 3402, 1360, 561 and 1110. Data are mean ± SEM. \**P* ≤ 0.05, \*\**P* ≤ 0.01, \*\*\**P* ≤ 0.001, \*\*\*\**P* ≤ 0.0001, unpaired Student’s *t*-test. **G-N,** Percentage of subtracks having directed motion **(G & K)**, normal diffusion **(H & L)**, subdiffusion **(I & M)** and confined diffusion **(J & N)** in WT, *MIC13* KO Hela cells expressing MIC60- and MIC10-SNAP and in WT HAP1 cells expressing MIC60- and MIC10-SNAP, *MIC60* KO HAP1 cells expressing MIC10-SNAP and *MIC10* KO HAP1 cells expressing MIC60-SNAP, respectively. Data are mean ± SEM. \**P* ≤ 0.05, \*\**P* ≤ 0.01, \*\*\**P* ≤ 0.001, \*\*\*\**P* ≤ 0.0001, unpaired Student’s *t*-test.

**Figure EV4.**
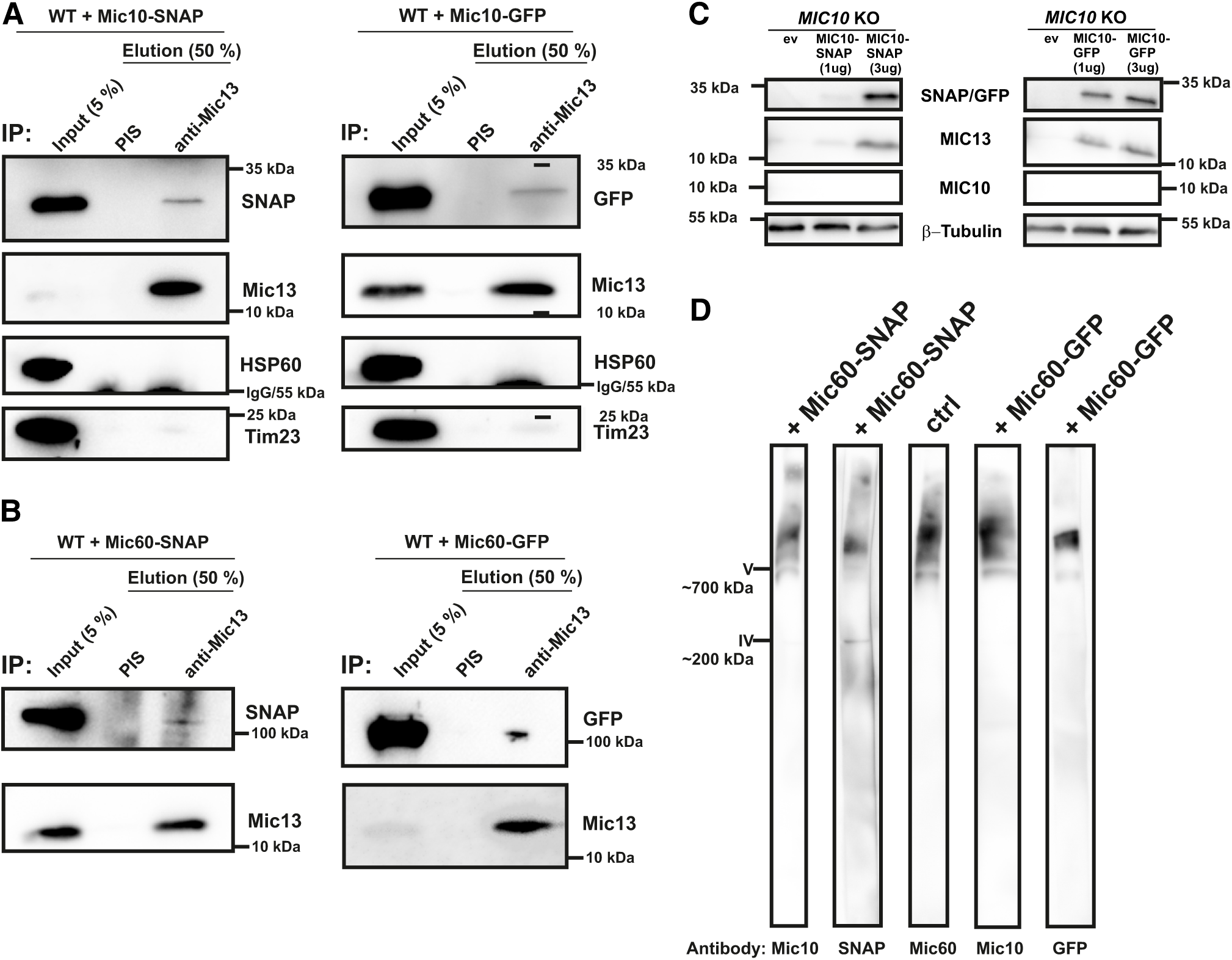
Functionality and localization of MIC10 and MIC60 is not impaired by C-terminal protein tags. **A, B** HEK293 cells expressing either MIC10-SNAP/GFP **(A)** or MIC60-SNAP/GFP **(B)** were used for coimmunoprecipitation using anti-MIC13. Endogenous MIC13 could pull down MIC10-SNAP or MIC10-GFP (A) as well as MIC60-SNAP or MIC60-GFP (B). **C** *MIC10* KO HAP1 cells expressing an empty vector (ev), 1 and 3 µg of MIC10-SNAP/GFP and probed with antibodies against SNAP/GFP, MIC10 and MIC13. MIC13 protein levels in *MIC10* KO HAP1 cells are rescued by expression of MIC10-SNAP (3 µg) and MIC10-GFP (1 and 3 µg) showing that the GFP and SNAP tags do not interfere with the MIC10 function. **D** The isolated mitochondria expressing MIC60-SNAP or MIC60-GFP were probed for their incorporation into the native MICOS complex. The complex containing MIC60-GFP or MIC60-SNAP runs at similar molecular weight as MICOS complex in native state marked by MIC60 or MIC10 antibody, confirming their incorporation into native complex.

## Movies Legends

**Movie 1 - MIC10-SNAP punctae marking CJs move dynamically within seconds**

Representative STED super-resolution time-lapse movie of Hela cell expressing MIC10-SNAP shows rapid movement of MIC10-SNAP punctae marking CJs. Box marks selection shown as a zoom in Movie 3.

**Movie 2 - MIC60-SNAP punctae marking CJs moves dynamically within seconds**

Representative STED super-resolution time-lapse movie of Hela cell expressing MIC60-SNAP shows rapid movement of MIC60-SNAP punctae marking CJs. Box marks selection shown as a zoom in Movie 4.

**Movie 3 - CJs marked by MIC10-SNAP undergo joining and splitting events**

Representative STED super-resolution time-lapse movie showing a magnified view of a mitochondrion expressing MIC10-SNAP marking CJs. Two MIC10 punctae rhythmically join and split as also shown in in Fig 4B. Green and magenta asterisks mark joining and splitting events respectively.

**Movie 4 - CJs marked by MIC60-SNAP undergo joining and spliting events**

Representative STED super-resolution time-lapse movie showing a magnified view of a mitochondrion expressing MIC60-SNAP, marking CJs. This movie is represented in Fig 4D. Green and magenta asterisks mark joining and splitting events respectively.

**Movie 5 - Joining and splitting of MIC60-SNAP punctae occurs in a MIC13-dependent manner**

Representative STED super-resolution time-lapse movie showing a magnified view of a mitochondrion from *MIC13* KO cells expressing MIC60-SNAP. This movie is represented in Fig 5B. Number of joining and splitting events are drastically reduced. Green asterisks mark a rare joining event.

**Movie 6 - MIC13-SNAP show that CJs and cristae undergo remodelling within seconds**

Representative STED super-resolution time-lapse movie, (2.5 s/frame) of Hela cell expressing MIC13-SNAP, shows rapid movement of CJs as well as cristae marked by MIC13-SNAP. Box marks selection shown as a zoom in Movie 7.

**Movie 7 - MIC13-SNAP show continuous cristae remodelling in form of fusion and fission cycles linked to CJs dynamics**

Representative STED super-resolution time-lapse movie showing a magnified view of a mitochondrion expressing MIC13-SNAP at a framerate of 2.5 s/frame. This movie is represented in Fig 6B. Various cristae remodelling events are marked by green and magenta asterisks representing fusion and fission of cristae respectively. Cristae remodelling event that forms a structure resembling letter ‘X’ is also marked.

**Movie 8 - MIC13-SNAP show that CJs and cristae undergo remodelling within seconds**

Representative STED super-resolution time-lapse movie (1.3 s/frame) of Hela cell expressing MIC13-SNAP shows rapid movement of CJs as well as cristae marked by MIC13-SNAP. Box marks selection shown as a zoom in Movie 9.

**Movie 9 - MIC13-SNAP show continuous cristae remodelling in form of fusion and fission cycles linked to CJs dynamics**

Representative STED super-resolution time-lapse movie showing a magnified view of a mitochondrion expressing MIC13-SNAP at a frame rate of 1.3 s/frame. This movie is represented in Fig 6D. Various cristae remodelling events are marked by green and magenta asterisks representing fusion and fission of cristae respectively.

**Movie 10 - ATP5I-SNAP marking cristae show cristae remodelling within seconds**

Representative STED super-resolution time-lapse movie (2.5 s/frame) of Hela cell expressing ATP5I-SNAP, marking cristae as transverse bridge across mitochondria, shows rapid movement of cristae. Box marks selection shown as a zoom in Movie 11.

**Movie 11 - ATP5I-SNAP marking cristae show continuous cristae remodelling in form of fusion and fission cycles**

Representative STED super-resolution time-lapse movie showing a magnified view of a mitochondrion expressing ATP5I-SNAP at a frame rate of 2.5 s/frame. This movie is represented in Fig 7B. Various cristae remodelling events are marked by green and magenta asterisks representing fusion and fission of cristae. A cristae remodelling event that forms a structure resembling letter ‘’ is also marked.

**Movie 12 - COX8A-SNAP marking cristae show continuous cristae remodelling in form of fusion and fission cycles**

Representative STED super-resolution time-lapse movie showing a magnified view of a mitochondrion expressing COX8A-SNAP at the frame rate of 2.6 s/frame. This movie is represented in Fig 7D. Various cristae remodelling events are marked by green and magenta asterisks representing fusion and fission of cristae.

**Movie 13 - Fusion and fission cycles of cristae marked by COX8A-SNAP are reduced in *MIC13* KO cells**

Representative STED super-resolution time-lapse movie showing a magnified view of a mitochondrion expressing COX8A-SNAP in *MIC13* KO cells at a frame rate of 2.6 s/frame. This movie is represented in Fig 7F. The number of fusion and fission events of cristae are reduced. Some rare events are marked by green and magenta asterisks representing fusion and fission of cristae.

**Movie 14 - Cristae dynamics helps redistribute membrane potential, marked by TMRM, between cristae**

Representative STED super-resolution time-lapse movie showing a magnified view of a mitochondrion stained with membrane potential specific dye TMRM at a frame rate of 1.5 s/frame. This movie is represented in Fig 8B. The green asterisks show the cristae which undergo fusion accompanied by immediate distribution of TMRM signal.

